# Neuromuscular embodiment of feedback control elements in *Drosophila* flight

**DOI:** 10.1101/2022.02.22.481344

**Authors:** Samuel C. Whitehead, Sofia Leone, Theodore Lindsay, Matthew R. Meiselman, Noah Cowan, Michael Dickinson, Nilay Yapici, David L. Stern, Troy Shirangi, Itai Cohen

## Abstract

While insects like *Drosophila* are flying, aerodynamic instabilities require that they make millisecond-timescale adjustments to their wing motion to stay aloft and on course. These stabilization reflexes can be modeled as a proportional-integral (PI) controller; however, it is unclear how such control might be instantiated in insects at the level of muscles and neurons. Here, we show that the b1 and b2 motor units—prominent components of the fly’s steering muscles system—modulate specific elements of the PI controller: the angular displacement (integral, I) and angular velocity (proportional, P), respectively. Moreover, these effects are observed only during the stabilization of pitch. Our results provide evidence for an organizational principle in which each muscle contributes to a specific functional role in flight control, a finding that highlights the power of using top-down behavioral modeling to guide bottom-up cellular manipulation studies.

---

To maintain stability, locomoting animals continuously update their motor actions based on sensory information^1, 2^. Such motor corrections are particularly important during extreme forms of locomotion like insect flight, where aerodynamic instabilities emerge rapidly when left uncorrected^3–5^. To contend with such instabilities, insects like *Drosophila* sense changes in their body orientation and respond with subtle modulations in wing motion on millisecond timescales^6–9^. This feedback control underlying *Drosophila* flight can be modeled by a set of proportional-integral (PI) controllers that describe the stabilization of all three rotational degrees of freedom: yaw^10^, pitch^11, 12^, and roll^13^. These PI controller models linearly combine the fly’s body angular velocity (P) and its angular displacement (I) to quantitatively predict changes in wing motion that counteract perturbations. Here, we use this powerful top-down framework for describing behavior to elucidate the bottom-up neuromuscular implementation of these flight stabilization reflexes.

*Drosophila* implement fast flight control—along with a suite of other impressive aerial maneuvers—using a sparse motor system consisting of 12 pairs of wing steering muscles (Fig. 1A)^14, 15^, each innervated by a single excitatory motoneuron^16^. Two of these 12 steering muscles that are thought to play a prominent role in flight control are the first and second basalar muscles, b1 and b2 (Fig. 1A)^15, 17–20^. These muscles regulate wing motion via their agonistic actions on the basalar sclerite, a skeletal element at the base of the wing^14^. Studies in *Drosophila* and *Calliphora* (blow flies) demonstrate that changes in either b1 or b2 muscle activity contribute to the modulation of wing stroke amplitude,^15, 17–21^ a primary control parameter used to stabilize both roll^13^ and pitch orientation^12^. Despite their similar effects on wing kinematics, however, the b1 and b2 muscles differ drastically in their physiology: b1 is tonically active during flight and encodes changes in wing kinematics via phase shifts in firing, whereas b2 is phasically activated during maneuvers, but generally quiescent during straight flight bouts^15, 17, 18^. Taken together, these studies suggest that b1 and b2 are both poised to play critical, but potentially distinct, roles in rapid flight control.

**Figure 1.**
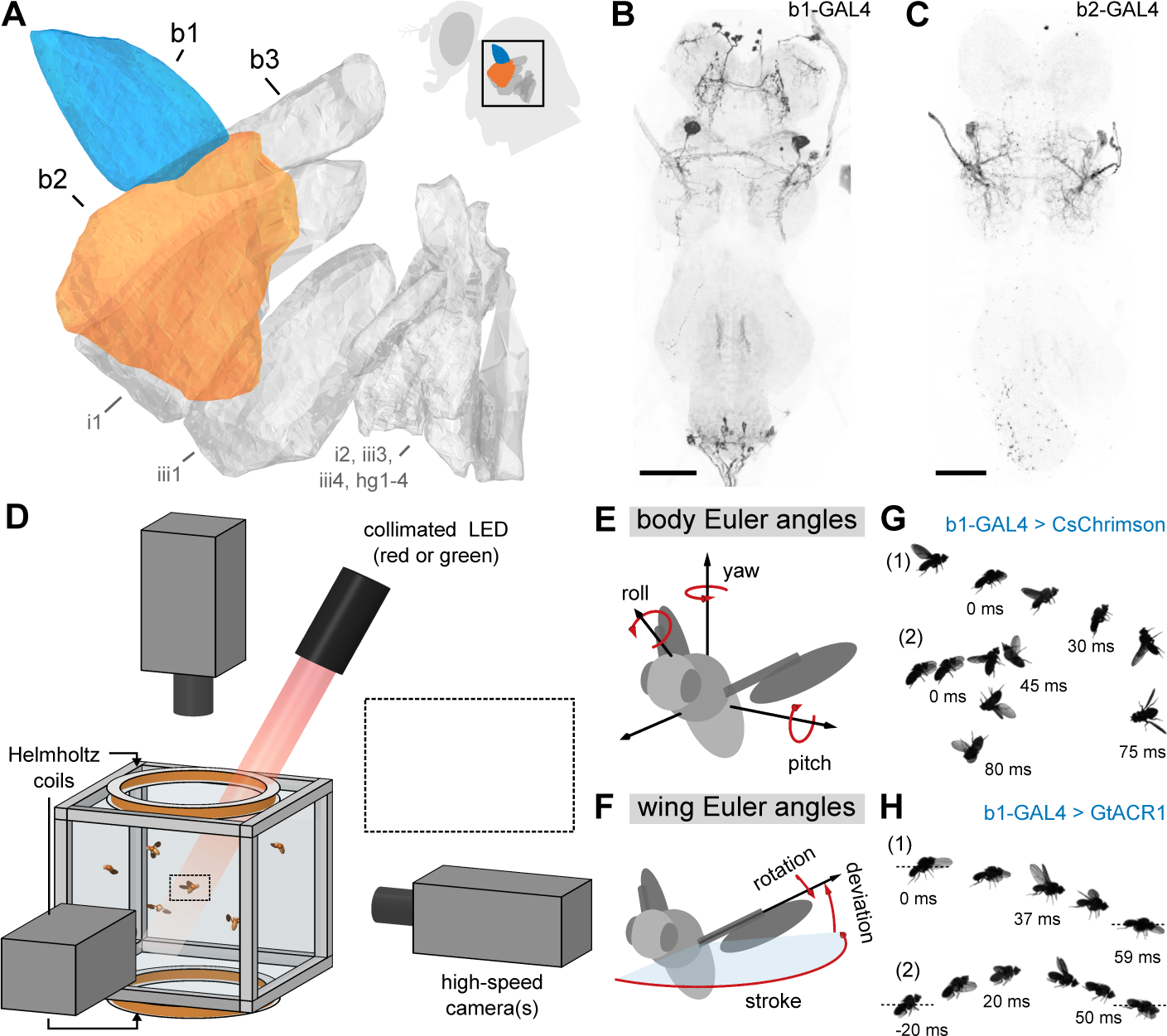
Combining genetic tools and free-flight apparatus to quantify effects of b1 and b2 manipulations. **A** Direct steering muscles of the *Drosophila* wing motor, with the first and second basalars (b1 and b2) highlighted in blue and orange, respectively. Data from^15^. Inset shows a fly silhouette, with a black box indicating the approximate position of the steering muscles. **B**,**C** Maximum intensity projections of the fly ventral nerve cord (VNC) expressing CsChrimson-mVenus (black) driven by *b1-GAL4* (**B**) and *b2-GAL4* (**C**). Light gray shows DNCad (neuropil). Scale bars 50 μm. **D** Schematic of experimental apparatus used to deliver optogenetic and/or mechanical perturbations to freely flying *Drosophila* while filming their maneuvers at 8000 frames*/*s. Inset illustrates magnetic field from Helmholtz coils interacting with magnetic pin glued to a fly to produce a perturbing pitch torque. Optical trigger (not shown) allows automated capture of hundreds of movies per trial. **E**,**F** Definitions of the body (**D**) and wing (**E**) Euler angles used to describe flight kinematics. **G**,**H** Photomontages of example flies undergoing optogenetic excitation (**G**; Mov. S1,S2) and silencing (**H**; Mov. S3,S4) of the b1 motoneuron using CsChrimson and GtACR1, respectively. Each panel shows photomontages of two flies—labeled “(1)” and “(2)”—with time stamps indicating timing relative to the onset of the 50 ms LED stimulus (*t*=0 ms). Full fly genotypes given in Table S2.

To first resolve the effects of b1 and b2 manipulation on free flight kinematics, we measured changes in wing and body dynamics of flies experiencing brief bouts of mid-air optogenetic excitation or inhibition (Fig. 1; Materials and Methods). In these experiments, we targeted the motoneurons of the b1 and b2 muscles using the split-GAL4 driver lines *MB258C-GAL4* (*b1-GAL4*; Fig. 1B, S1A–B; Tab. S1) and *b2-SG*^22^ (*b2-GAL4*; Fig. 1C, S1C–D; Tab. S1) to drive expression of CsChrimson^23^ (excitation) or GtACR1^24^ (inhibition). Using the flight chamber shown in Figure 1D, we captured and quantified flight kinematics (Fig. 1E,F) prior to, during, and after the application of a 50 ms light pulse. Figure 1G,H show photomontages of responses to b1 motoneuron excitation (G) and inhibition (H) viewed from the side. As illustrated in Figure 1G, excitation of the b1 motoneuron evoked extreme upward pitching maneuvers, with the fly rotating ≥90° during the 50 ms period of stimulation (Mov. S1, S2). Under b1 motoneuron inhibition, flies pitched downwards, dipping to angles below the horizontal plane (Fig. 1H; Mov. S3, S4)).

To quantify these kinematic changes, we analyzed hundreds of such flight videos (Fig. 2). Optogenetic excitation of both *b1-GAL4* and *b2-GAL4* flies drove large, nose-up deviations in pitch, with smaller deviations in roll and yaw (Fig. 2A, blue, orange; Mov. S5,S6). These changes in pitch orientation were driven by bilateral modulations of wingbeat angles during stimulation (Fig. 2B). During stimulated wingbeats, we observed an increase in the forward stroke angle (Fig. 2C, top), and a corresponding increase in the aerodynamic pitch torque, estimated using a quasi-steady model^25^ (Fig. 2C, bottom; Supplementary Text). In contrast, genetic control experiments using the empty split-GAL4 line *SS01062-GAL4*^26^ (aka *empty*) with the *UAS-CsChrimson* transgene showed no measurable changes in body orientation or wing motion upon optogenetic excitation (Fig. 2A–C, gray; Mov. S7). Overall, our findings are consistent with previous electrophysiological studies in flies, which showed that increased activity in either b1 or b2 drove marked increases in wingstroke parameters like the downstroke deviation and forward stroke angles^17–20^ (Fig. S2).

**Figure 2.**
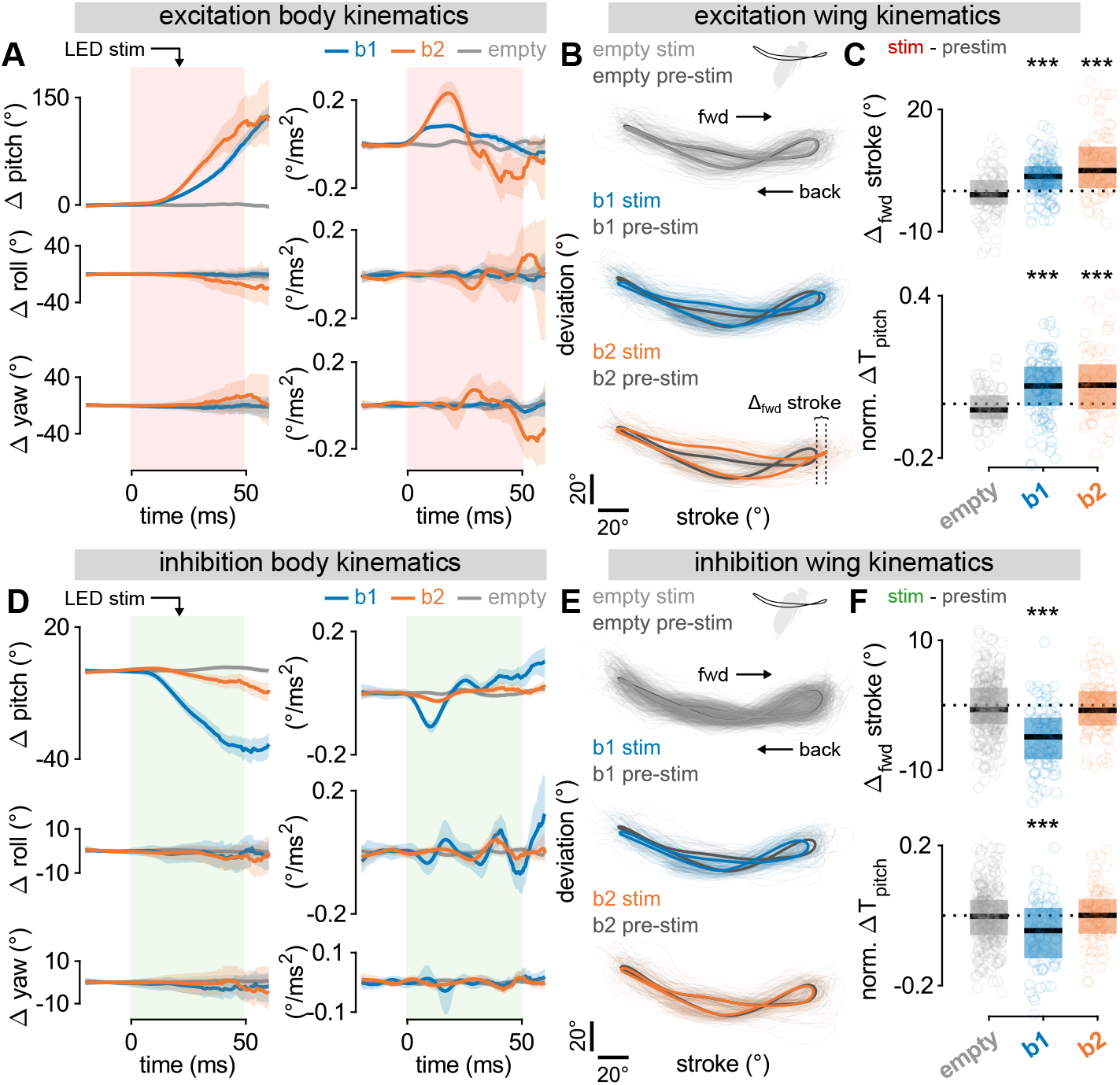
In-flight optogenetic activation and silencing of the b1 and b2 motoneurons drive changes to pitch orientation. **A** Body kinematics versus time in response to 50 ms optogenetic activation of *b1-GAL4* (blue; *N*=140 movies), *b2-GAL4* (orange; *N*=84 movies), and *SS01062-GAL4* (aka *empty*; gray; *N*=108 movies) flies with CsChrimson. Rows correspond to rotational degrees of freedom: pitch (top), roll (middle), and yaw (bottom). Columns give angular displacement (left) and angular acceleration (right). Data shown represent mean ± 95% confidence interval. **B** Wing kinematic data for movies in **A**. Plots show wing tip angular position in the wingstrokes prior to (dark gray; “pre-stim”) and during (light gray, blue, orange; “stim”) optogenetic activation. Thick traces represent population averages; thin lines represent single-fly wingbeats. Vertical and horizontal scale bars provide 20° references for deviation and stroke angles, respectively. **C** Change in forward stroke angle (top) and normalized, wingbeat-averaged aerodynamic pitch torque (bottom) for wingbeats prior to and during optogenetic activation of *b1-GAL4* (blue), *b2-GAL4* (orange), and *SS01062-GAL4* (gray) flies. Circles show raw data; box and horizontal line show interquartile range and median, respectively. Statistical significance determined via Wilcoxon signed-rank test (***, *p*<0.001; **, *p*<0.01; *, *p*<0.05). **D**–**F** Same as in **A**–**C**, but with optogenetic silencing of *b1-GAL4* (blue; *N*=89 movies), *b2-GAL4* (orange; *N*=89 movies) and *SS01062-GAL4* (gray, *N*=323 movies) flies with GtACR1. Full fly genotypes given in Table S2.

Performing the same analyses with optogenetic silencing, we found that b1-silenced flies primarily underwent large, nose-down changes to pitch, while b2-silenced and genetic control flies showed little change to their body kinematics (Fig. 2B, blue, orange; Mov. S8–S10). Correspondingly, we observed the largest change in b1-silenced flies’ wingstrokes (Fig. 2E, blue), with a significant decrease in both forward stroke angle and resulting pitch torque as compared to both b2-silenced and the genetic control flies (Fig. 2F). This lack of response in b2-silenced flies is consistent with the fact that b2 is a phasic muscle^15^, and would thus likely be quiescent during steady-state, unperturbed flight. Collectively, these results indicate that changes in bilateral b1 motoneuron activity are capable of bidirectionally modulating pitch torque, whereas bilateral manipulation of the b2 motoneuron activity results only in pitch up torque. This ability to affect pitch orientation confirms that both muscles could play an important role in the control of this degree of freedom.

To directly test the contributions of these muscles to flight stabilization, we quantified responses to imposed mid-air perturbations from flies with chronically inhibited b1 and b2 activity. To conduct these experiments, we drove expression of the inwardly rectifying potassium channel Kir2.1^27, 28^ in the b1 and b2 motoneurons (Tab. S1, Materials and Methods). Despite the kinematic responses observed in optogenetically b1-silenced flies (Fig. 2D–F), chronic silencing of the b1 motoneuron (Fig. S4) did not preclude flight, a phenomenon we suspect can be attributed to compensatory activity in other wing steering muscles. To assay the effects of this chronic silencing on stabilization maneuvers, we imposed rapid, mid-air magnetic perturbations to freely flying flies with magnetic pins glued to their backs (Fig. 3A, S3A). Using custom tracking software^29^, we extracted the flies’ corrective wing and body kinematics as they responded to either pitch or roll perturbations.

**Figure 3.**
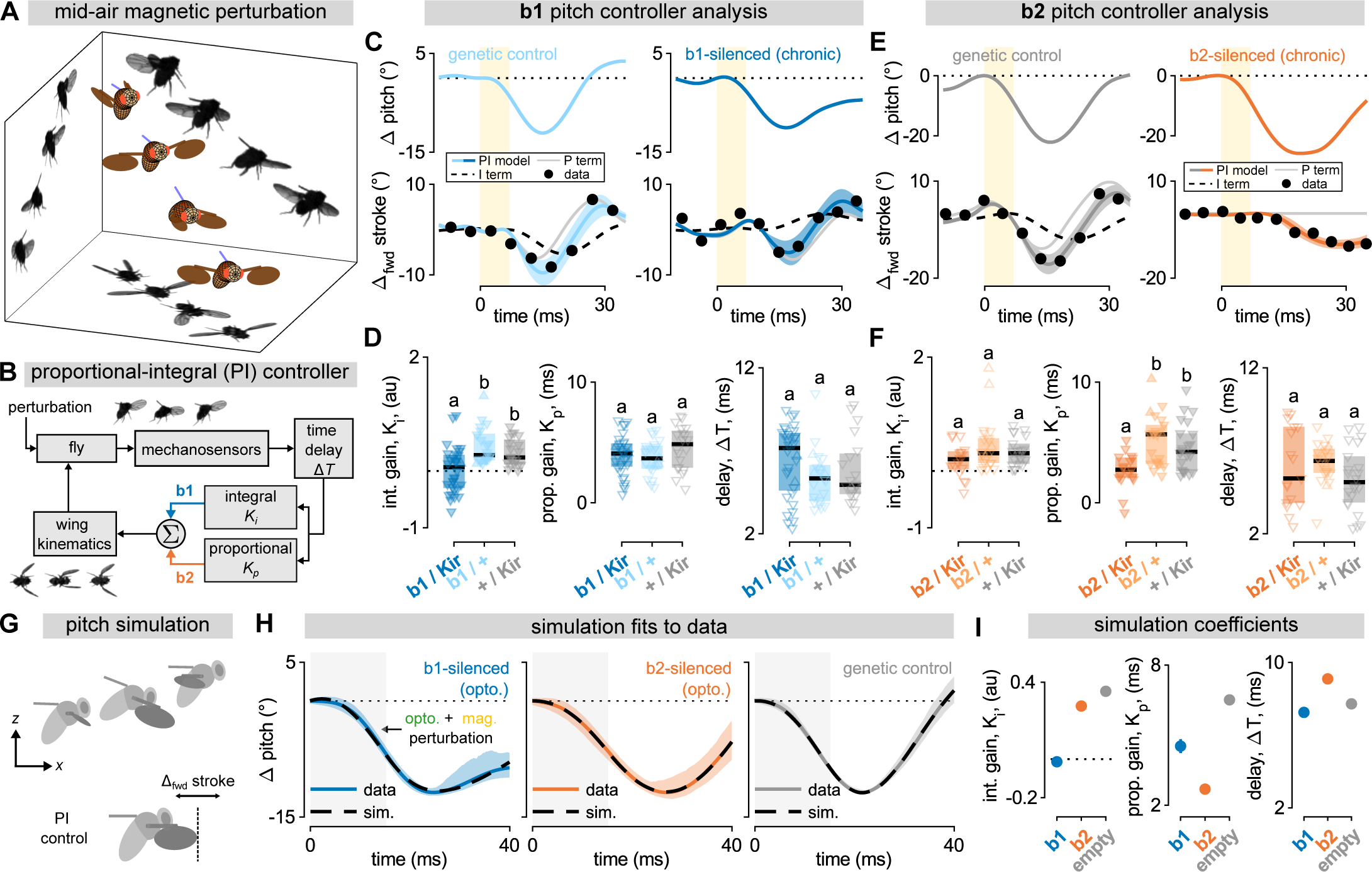
Inhibiting the b1 and b2 motoneurons alters the integral and proportional gains for pitch control, respectively. **A** Reconstruction of a fly experiencing and correcting for a pitch down mechanical perturbation. The walls show photomontages from three high-speed cameras, with the 3D rendered fly showing measured kinematics. Attached ferromagnetic pin false-colored blue. **B** Proportional-integral (PI) controller model for rapid flight stabilization, as in Equation 1. External perturbations and/or wing kinematic changes alter the fly’s dynamics; the fly measures the resultant changes to angular velocity using mechanosensory organs. This angular velocity signal is subject to a time delay, and then split into proportional and integral branches, the weighted sum of which determines subsequent changes to wing kinematics, producing a corrective counter torque. Blue and orange arrows highlight experimental findings showing that inhibition of the b1 and b2 motor units attenuates the integral and proportional feedback responses, respectively. **C** Example pitch down perturbation analyses for a genetic control fly (left; Mov. S11) and a b1-silenced fly (right; Mov. S12). Top row shows change in body pitch angle over time (blue traces); bottom row shows measured change in forward stroke angle over time (black dots), PI controller model fit to data (blue traces with 95% CI), P term (proportional; thin gray line), and I term (integral; dashed black line). Yellow bar indicates 7 ms magnetic pulse. **D** Summary statistics for PI controller model parameters from Equation 1—integral gain (*K*_i_; left), proportional gain (*K*_p_; center), and time delay (Δ*T*, right)—for b1-silenced flies (dark blue; *N*=32) and two genetic controls (light blue and gray; *N*=21 and 20). Each data point represents a pitch perturbation movie. Upward and downward triangles represent pitch up and down perturbations, respectively. Box plots show median (black line) and interquartile range for each genotype. Lower case letters (i.e. “a”,”b”) above data indicate significance categories, determined via Wilcoxon rank sum test with Bonferroni correction (*α*=0.05). **E** Same as in **C** but with a genetic control (left; Mov. S13) and b2-silenced fly (right; Mov. S14). The integral term (dashed black line) is not visible in the bottom right panel because it is covered by the full PI controller fit (solid orange line). **F** Same as **D** but with b2-silenced flies (dark orange; *N*=17) and two genetic controls (light orange and gray; *N*=22 and 22). **G** Illustration of time snapshots from longitudinal flight simulation (top) and change in forward stroke angle as a control parameter according to Equation 1 (bottom). **H** Change in body pitch angle over time for simulated flies (dashed black lines) fit to experimental data (population average with 95% CI envelope). Columns correspond to different genotypes: *b1-GAL4 > GtACR1* (“b1-silenced”; left; *N*=41 movies), *b2-GAL4 > GtACR1* (“b2-silenced”; middle; *N*=32), and *empty > GtACR1* (“genetic control”; right; *N*=62). Gray bar represents the 15 ms simultaneous LED and magnetic field stimuli, which optogenetically silence and impose external torque, respectively. **I** PI controller parameters from simulation fits in **H**—integral gain (*K*_i_, left), proportional gain (*K*_p_, middle), and time delay (Δ*T*, right)—for each genotype. Error bars show 95% CI. Full fly genotypes given in Table S2.

The observed kinematic changes under b1 or b2 inhibition become particularly transparent in the context of a proportional-integral (PI) controller framework^10, 12, 13^ (Fig. 3B), which provides a reduced-order description for the corrective response. In the case of pitch perturbations^11, 12^, this PI model predicts changes in forward stroke amplitude (Δ_fwd_*ϕ*) as a function of time (*t*):

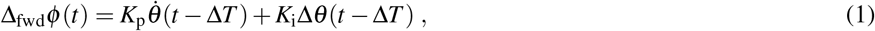

where *K*_p_, *K*_i_, and Δ*T* are the proportional gain, integral gain, and time delay of the PI controller, respectively. Thus, the change in forward stroke angle (Δ_fwd_*ϕ*) at time *t* is quantitatively predicted by a linear combination of the body pitch angular displacement (Δ*θ*) and pitch velocity 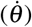 at an earlier time point, *t* − Δ*T*. The controller gain coefficients, *K*_p_ and *K*_i_, determine the relative weights of angular velocity and displacement to the corrective response; the time delay, Δ*T*, corresponds to the reflex latency. Comparing these controller parameters in b1- and b2-silenced flies versus genetic controls, allows for directly testing the roles of the b1 and b2 motor units in the reflex response.

We illustrate this strategy by comparing pitch perturbation events for individual flies: one from a genetic control group (Fig. 3C, left; Mov. S11), and one from the b1-silenced group (Fig. 3C, right; Mov. S12). For similar maximum pitch deflections, the control group fly was able to return to its original orientation roughly 25 ms after the onset of the magnetic field pulse, while the b1-silenced fly leveled off at a pitch angle below its pre-perturbation orientation (Fig. 3C, top). In both cases, the PI controller models (Fig. 3C, bottom) quantitatively predict the time course of forward stroke angle (Δ_fwd_*ϕ*); for the b1-silenced fly, however, the *integral* gain (*K*_i_) obtained from the fit is negative. This result is counterintuitive, as it would indicate a control law pushing the fly away from its initial orientation. Overall, b1-silenced flies showed a statistically significant decrease in the integral gain (*K*_i_) of the PI controller model as compared to genetic controls (Fig. 3D, left), whereas the proportional gain (*K*_p_) and time delay (Δ*T*) were not significantly different across genotypes (Fig. 3F, middle and right). Performing the same analysis on roll stabilization, we found no effect of b1 motoneuron silencing on feedback control (Fig. S3). Taken together, these results indicate that b1 motoneuron silencing primarily affects pitch stabilization, and does so in a manner that is captured by a single parameter in the PI controller model: the integral gain, *K*_i_ (Fig. 3B; blue arrow).

We applied the same strategy to elucidate the role of b2 in the stabilization reflexes (Fig. 3E,F). Pitch perturbation events for a genetic control fly (Fig. 3E, left; Mov. S13) and a b2-silenced fly (Fig. 3E, right; Mov. S14) are once again well fit by the PI controller model. Here, however, the PI controller fit to the corrective response of the b2-silenced fly lacks a *proportional* term, i.e. *K*_p_ = 0. This trend holds across flies: compared to genetic controls, b2-silenced flies exhibited reduced proportional gain, whereas the distributions of other controller coefficients (*K*_i_and Δ*T*) were not significantly different (Fig. 3F). We performed the same analyses using roll perturbations, and found that b2 silencing had no measurable effect on roll stabilization (Fig. S3). Thus, silencing the b2 motoneuron uniquely affected the proportional term, *K*_p_, for pitch control (Fig. 3B; orange arrow).

This simple interpretation—b1 and b2 actuating integral and proportional control for pitch stabilization, respectively—is rein-forced by matching simulations of flapping flight^3, 12^ to experimental data from optogenetically silenced flies undergoing magnetic perturbations (Fig. 3G–I; Supplementary Text). Here, we optimized the PI controller parameters in simulated flies to best reproduce measured kinematics in real flies. Using this approach, we quantitatively captured the changes in body pitch angle over time for flies undergoing simultaneous magnetic perturbation and optogenetic silencing (Fig. 3H). Consistent with our previous results, the controller parameters (*K*_i_, *K*_p_, and Δ*T*) obtained from these simulation fits show that b1 silencing reduces integral gain (Fig. 3I; left), while b2 silencing reduces proportional gain (Fig. 3I; middle).

Our finding that the b1 and b2 motor units act as elemental control features in the pitch flight controller of *Drosophila* confirms a previous hypothesis that the two physiological categories of steering muscles—tonic (e.g. b1) and phasic (e.g. b2)—respectively actuate integral and proportional control^15^. Based on these results and previous studies showing a correspondence between anatomical groupings and their recruitment for maneuvers about different rotational axes^14, 15^, we conjecture that other tonic and phasic muscles in the wing motor system might be similarly mapped onto the integral and proportional controller parameters for yaw and roll. Importantly, testing this conjecture will once again require combining top-down behavioral modeling in freely flying animals and bottom-up manipulations for probing neuromuscular systems^2, 30^, as demonstrated here. Such hybrid experimental approaches have the potential to become even more powerful as genetic tools for precise, cell-specific interventions continue to be developed. In combination with electron microscopy connectomics^31^, this experimental paradigm will likely provide critical insights into the full sensorimotor cascade for the PI controller, as well as other motor control circuits being identified.

## Acknowledgements

We thank Tsevi Beatus for advice on data analysis techniques; Anne von Phillipsborn, Yoshi Aso, Gwyneth Card, Wyatt Korff, and Erica Ehrhardt for providing reagents; Jesse Goldberg and Joe Fetcho for guidance and support; Meera Ramaswamy for general feedback; and the members of the Cohen Lab for myriad useful conversations. Further, we are grateful to the Biotechnology Resource Center (BRC) at Cornell for the use of their imaging tools, as well as the Cornell Statistical Consulting Unit (CSCU)—particularly Lynn Johnson—for assistance with statistical methods.

This work was supported by:

National Institute of Health (NINDS, 1R01NS116595)

National Science Foundation (IOS, 1546710; IOS 1452510 (to M.H.D))

Janelia Research Campus Visitor Project Program

Department of Defense (NDSEG Fellowship)

Army Research Office (61651-EG)

Office of Naval Research (N00014-21-1-2431 (to NJC))

## Author contributions statement

Conceptualization: SCW, IC, TS, DS

Methodology: SCW, IC, TS, DS, NY, TL,

MD Resources: SCW, IC, NY, TS, DS, MD

Investigation: SCW, TL, SL, MM

Formal Analysis: SCW, NC Software: SCW

Visualization: SCW

Writing – original draft: SCW, IC

Writing – reviewing and editing: SCW, IC, TS, DS, MD, NC, MM, NY

## Additional information

### Competing interests

The authors declare that they have no financial or other competing interests.

### Data and materials availability

Processed data and code will be available through Cornell University Library’s institutional repository, eCommons. Raw high-speed video data is available upon request.

## Materials and Methods

### Fly stocks and fly handling

Flies used for optogenetics experiments reared in the dark at room temperature on 0.4 mM retinal food (Media Facility, Janelia Research Campus). Flies used for all other experiments (e.g. mechanical perturbation) were raised at room temperature on standard fly media made from yeast, agar, and sucrose with a 12 h light/dark cycle. Female flies, 3-6 days post eclosion, were used for all flight experiments. A full list of *Drosophila melanogaster* stocks used in this paper is given in Table S1.

### Immunohistochemistry

Light microscopy images in Figures 1, S1, and S5 were obtained using a protocol similar to the one described in^1^. In brief, full central nervous systems were dissected into PBS (phosphate-buffered saline) and fixed in 4% paraformaldehyde in PBS for 35 minutes at room temperature. Fixed tissues were then washed in PBT (PBS containing 0.1% Triton X-100) and incubated with primary antibodies diluted in PBT overnight at 4 ^*°*^C. The next day, samples were washed in PBT for several hours at room temperature and then incubated with secondary antibodies diluted in PBT overnight at 4 ^*°*^C. Samples were then washed all day with PBT, placed onto polylysine-coated coverslips, dehydrated through an ethanol series, cleared in xylenes, and mounted in DPX (Sigma-Aldrich). Adult CNS tissues were then imaged on a Leica SP6 confocal microscope with optical sections at 0.3 mm intervals. Maximum intensity projections (as shown in Figures 1B,C; S1A–D; and S5A,B) were generated using ImageJ.

Phalloidin images were obtained using a protocol similar to the one detailed in^2^. In brief, a razor blade was used to hemisect thoraces of adult female flies frozen in Tissue-Tek O.C.T. (electron Microscopy Sciences, Cat#62550-01). Samples were then fixed in a solution of 4% paraformaldehyde in PBS for 45 minutes and subsequently washed three times in PBT. Primary antibodies and phalloidin stain were then added, and the samples were nutated for 7-10 days at 4 ^*°*^C. After the staining period, samples were rinsed in PBT and cleared using the SeeDB protocol^3^. Samples were then mounted with SeeDB in between two glass coverslips, with another glass coverslip placed on top and clear nail polish used to seal the sample in. These samples were subsequently imaged using a Zeiss LSM 880 upright confocal microscope.

The following stains/antibodies were used in the above protocols: rabbit polyclonal anti-GFP (ThermoFisher, Cat#A11122), rat anti-DN-cadherin (DSHB, DN-Ex #8), Alexa Fluor Plus 405 Phalloidin (Invitrogen, Cat#A30104), rabbit polyclonal anti-GFP (Torrey Pines, Cat#TP401), and Alexa Fluor 488 goat anti-rabbit (Invitrogen, Cat#A27034).

### Fly preparation

For perturbation experiments, individual flies were anesthetized at 0 ^*°*^C–4 ^*°*^C, at which point we carefully glued 1.5–2 mm long, 0.15 mm diameter ferromagnetic pins to their notum (dorsal thoracic surface). The pins were oriented to lie in the fly’s sagittal plane for pitch perturbation experiments; for roll experiments the pins were oriented perpendicular the fly’s sagittal plane. Experiments with unpinned flies showed that the addition of the pin did not qualitatively alter flies’ flight kinematics. The attachment of pins adds mass that is comparable to natural intra-fly mass variation, and adds negligibly to the off-diagonal components of the fly’s inertia tensor (for detailed calculations, see^4, 5^).

### High-speed videography

We performed experiments with 15–30 flies prepared as above, all with the same genotype. We released these flies into a transparent cubic flight chamber with side length 13 cm. The center of the chamber was filmed by three orthogonal high-speed cameras (Phantom V7.1) at 8000 frames*/*s and 512 × 512 pixel resolution, with the three cameras sharing a mutual filming volume of *∼*8 cm^3^ (Fig. 1D). Each camera was backlit by a focused 850(30) nm near-infrared LED (Osram Platinum Dragon). An optical trigger—created using split, expanded beams from a 5 mW, 633 nm HeNe laser (Thorlabs, HRR050) passed through a neutral density filter (Thorlabs, NE20A) with optical density 2.0, and incident upon two photodiodes (Thorlabs, FDS100)—was used to detect the entrance of flies into the filming volume of the high-speed cameras during experiments and initiate filming^6^. Prior to each experiment, we calibrated the cameras using the *easyWand* system from^7^.

### Optogenetic experiments

For each optogenetic experiment, 10–30 flies were released into the flight chamber described above for approximately 12 hours. To apply mid-air optogenetic stimulation, we used the optical trigger circuit described above to deliver a 50 ms bout of light stimulation from a collimated LED source placed outside the chamber (Fig. 1D) whenever a flying fly entered the center of the filming volume. This trigger also initiated filming with the high-speed cameras, which recorded flight activity prior to, during, and after the application of the light stimulus at 8000 frames*/*s. To apply the light stimulus, the optical trigger circuit drove a 50 ms duration voltage pulse to a LED driver (Thorlabs, LEDD1B), which was connected to either a 625 nm red LED (Thorlabs M625L4) or a 565 nm green LED (Thorlabs M565L3) for optogenetic excitation (CsChrimson) or inhibition (GtACR1) experiments, respectively. Both red and green LED sources were outfitted with a collimating attachment (Thorlabs, COP2-A) to generate a 50 mm diameter beam profile. The cross-sectional area of this beam was large enough so that a fly anywhere in the filming volume of all three cameras would necessarily be hit by the light source, and the collimation ensured that the stimulus intensity was uniform regardless of the fly’s location within the filming volume.

Unless otherwise noted, the stimulation LEDs were driven with a 1 A current, resulting in intensities of 731 μW*/*mm^2^ and 316 μW*/*mm^2^ for the red and green LEDs, respectively. Despite the optical trigger’s 633 nm HeNe laser ostensibly falling in the range of CsChrimson sensitivity, the optical filters on this light source ensured that the laser’s intensity, *∼*0.16 μW*/*mm^2^, was 2–3 orders of magnitude lower than any applied LED stimulus. Moreover, we did not observe any changes to flight behavior resulting from the HeNe light when LED stimulation was withheld.

To prevent outside light contamination during these these optogenetic experiments, the entire flight apparatus was surrounded by blackout curtains. Because flies are unlikely to initiate flight bouts in total darkness, a dim, blue fluorescent light bulb was used to illuminate the arena during experimental trials.

### Mechanical perturbation experiments

For each mechanical perturbation experiment, 10–20 flies were prepared by gluing small ferromagnetic pins to the dorsal side of their thoraces (see above) and subsequently released into the flight chamber. Similar to the optogenetic experiments described above, an optical trigger circuit was used to apply a variable-duration magnetic field pulse whenever a flying fly entered the center of the filming volume. High-speed cameras were used to record flight activity prior to, during, and after the application of the magnetic field at 8000 frames*/*s.

The impulsive magnetic field was generated by the optical trigger supplying a rapid current pulse to a pair of Helmholtz coils mounted on the top and bottom faces of the flight chamber. Due to the positioning of the Helmholtz coils, this produced a roughly uniform vertical magnetic field in the center of the filming volume, triggered by the entrance of a fly entered into this region of the flight chamber. Typical magnetic field strengths were on the order of *∼*10^*−*2^ T. The magnetic field from the coils acted on the magnetic moment of ferromagnetic pin glued to the fly, in turn generating a moment about either the fly’s pitch or roll axis, depending on the relative orientation of the field and pin (Fig. 1D, inset). Further details of this procedure are described in^4, 5, 8^.

Most experiments using this method for imposing external magnetic torques were performed with chronically silenced flies (Fig. 3C–F), with the magnetic field applied for 7 ms. However, this method could also be combined with the protocol for mid-air optogenetic manipulation described above in order to both optogenetically silence and mechanically perturb the same fly in a single movie. Combined optogenetic and mechanical perturbations were performed in two ways. In the first, the LED and Helmholtz coils were powered simultaneously (Fig. 3H,I) for 15 ms. In the second, the two signals were temporally offset, and given different durations (Fig. S7). Specifically, the LED was turned on for the time range *t* = 0 ms–50 ms, while the magnetic field was applied for *t* = 15 ms–22 ms, resulting in a 7 ms magnetic field pulse beginning 15 ms after the onset of the optogenetic LED (see Fig. S7 schematic).

### Flight data selection and kinematic extraction

Of the data collected in both optogenetic and mechanical perturbation experiments (as described above), we restricted our attention to videos that were amenable to kinematic analysis. Broadly, we required flight movies to contain the fly in the field of view of all three high-speed cameras long enough to analyze pre- and post-perturbation-onset flight kinematics. In our temporal coordinates, the perturbation onset occurs at time *t* = 0 ms, so we required the fly to be visible from all three camera views in the range *t* ∈ [*−*10 ms, 30 ms] for a particular movie to merit analysis. For just mechanical perturbation experiments, we imposed a slightly stricter set of criteria in addition to this time limit. Namely, we required that the perturbation act primarily along a single rotational axis and that there was no evidence of the fly performing a volitional maneuver prior to perturbation. Both of these criteria were imposed in an attempt to cleanly isolate corrective maneuvers for a single rotational degree of freedom.

To extract kinematic data from the three high-speed camera views, we used the custom-developed 3D hull reconstruction algorithm detailed in^6^. Using this algorithm, we obtained a 12 degree-of-freedom description of the fly—the 3D position of the fly center of mass and the three full sets of Euler angles for the fly body, left wing, and right wing—at each time point. For time points in which occlusion precluded the direct extraction of a particular kinematic variable, we used a cubic spline interpolant to fill in missing data values. For most analyses, raw body kinematics were filtered using a 100 Hz low-pass filter. Raw wing kinematics were smoothed using the Savitzky–Golay method. For the wing stroke angle, we used a polynomial order 7 with window size 21 frames (2.625 ms; for the wing deviation and rotation angles we used a polynomial order five and window size 11 frames (1.375 ms). Figure S8 shows example wing and body Euler angles from the perturbation movies shown in Figure 3C,E (Mov. S11, S12, S13, and S14) as an illustration of this process of kinematic extraction.

### Controller model fitting for single-trial data

For each movie of a fly performing a corrective maneuver in response to a mechanical perturbation—as in Figures 3C–F, S3C–F, S6, and S7—we fit a proportional integral (PI) model to the kinematic data obtained as above. The equations for pitch and roll PI controller models are given in Equations 1 and S1, and are based on previously derived models from^4, 5, 8^. To fit the controller coefficients, we performed a nonlinear least squares fit (Levenberg-Marquardt) for the gain coefficients, *K*_i_and *K*_p_, along with a grid search for the time delay, Δ*T*. Uncertainty in fit parameters was estimated using the fit parameter covariance matrix at the objective function minimum. The covariance matrix was approximated as *C* ≈ *σ* ^2^(*J J*)^*−*1^, where *C* is the covariance matrix, *σ* ^2^ is the variance of the fit residuals, and *J* is the calculated Jacobian at the objective function minimum.

### Fitting flight simulation parameters for averaged data

Our flapping flight simulation, similar to ones previously reported^5, 9^, poses the equations of motion for the fly in two translational (forward and vertical) and one rotational (pitch) degrees of freedom to study the dynamics of pitch stabilization in a reduced order framework. We used a set of analytic expressions^9^ to prescribe the wing motion of simulated flies, with changes to the wing forward stroke angle implemented continuously based on the PI controller scheme described above (Fig. 3; Eq. 1), which allowed us to calculate instantaneous aerodynamic forces and torques acting on the wings using a quasi-steady model^10, 11^. To mimic the details of our experiment, we included an option to impose an external pitch torque of tuneable magnitude and duration of the simulated fly. For each simulation run, we solved the equations of motion for the fly using MATLAB’s delayed differential equation solver, `dde23`. For more details on the flight simulation, see Supplementary Text.

When fitting simulation results to averaged experimental data we held all parameters but the three PI controller parameters (*K*_i_, *K*_p_, Δ*T*) and the strength of the perturbing pitch torque. The cost function minimized in each fit was the least squares difference between the simulated and measured body pitch angle in a 40 ms window beginning at the onset of the perturbation. We performed this non-linear least squares fit using the Levenberg-Marquardt algorithm in MATLAB’s `lsqnonlin` function, with ≥12 randomized start points to avoid the solver getting trapped in local minima. To characterize the uncertainty in the final fit parameters, we estimated 95% confidence intervals for each fit parameter using the procedure described above for controller model fits to data.

For the simulation fits shown in Figure 3H,I, we used data used from experiments in which flies expressing the optogenetic silencer GtACR1 under different GAL4 driver lines—*b1-GAL4, b2-GAL4*, and *SS01062-GAL4*—were subjected to simultaneous LED and magnetic field pulses lasting 15 ms, thereby transiently inhibiting the targeted neurons and applying a mechanical perturbation. To lessen the influence of motion/rotation about degrees of freedom other than pitch, we took population averages across genotypes as the basis for our simulation fits. Because the strength of the mechanical perturbation in our behavioral experiments varies depending on pin length, magnetization, and orientation, we first normalized the body pitch traces from each individual movie, so that each time series had identical maximum pitch deflections. We then averaged these normalized traces, and re-scaled the resulting average to have maximum pitch deflection equal to the median perturbation amplitude across all movies (*∼*12°). This ensured that the population-averaged traces for each genotype had matching perturbation magnitudes.

### LexA/Gal4 Intersectional Strategy

To account for the off-target expression in the brain of *b1-GAL4* flies (Fig. S5A), we used an intersectional approach to restrict Kir2.1 expression to the VNC in the b1 motoneuron chronic silencing experiments presented in Figure 3C,D. In this approach, akin to the one used in^12^, a Flp recombinase was driven by tshLexA (which expresses LexA in most neurons of the VNC) and used to excise a transcriptional stop cassette from a *10XUAS-Kir2*.*1* transgene, which in turn was driven by *b1-GAL4* (see Tab. S1).

As a control for the presence of additional transgenes introduced in this intersectional approach, we performed a set of pitch and roll perturbation experiments using *5XUAS-Kir2*.*1* crossed to *b1-GAL4* flies lacking the LexA/LexAop transgenes, a cross that mirrors the experiments in Figure 3E,F using *5XUAS-Kir2*.*1* flies crossed to *b2-GAL4* flies. The results of these experiments, presented in Figure S6, were consistent with those reported in the main text.

### Calcium imaging

To image calcium activity in steering muscles, we tethered flies in an upright orientation similar to free flight and mounted them in the custom imaging rig described in^2^. A 470 nm light-emitting diode (LED, Thorlabs) was used as an excitation light source and passed through a Chroma filter cube with a 480/40 nm excitation filter and 510 nm long-pass dichroic. GCaMP6f fluorescence was collected through a 10 *×* 0.45 numerical aperture (NA) lens and 535/50 nm emission filter. Stroke amplitude and wingbeat frequency were simultaneously monitored during fluorescence imaging: the former using a camera (Basilar) and infrared light, and the latter with a photodiode-based wingbeat analyzer.

We used the machine vision system Kinefly^13^ to extract stroke amplitude in real time, which we in turn used as feedback for closed-loop control of visual stimulus. This visual stimuli was displayed to the fly using a cylindrical panoramic display screen made from LED panels, with 470 nm peak wavelength, as in^2^. Using this, we presented the fly with both open-loop visual displays simulating rotation about the yaw, pitch, and roll axes, as well as closed-loop stripe fixation patterns.

From images of GCaMP6f fluorescence, we used the method of demixing described in^2^ to extract signals from individual muscles. In brief, this involved fitting measured signals to a generative model for muscle fluorescence based on anatomical priors and properties of the imaging apparatus. Since many of the muscles do not produce appreciable GCaMP6f signals during quiescence, we calculated the change in fluorescence, Δ*F/F*, with a baseline (*F*) determined by the first percentile of fluorescence signal on a per fly and trial basis.

## Supplementary Text

### Calcium imaging

To confirm that silencing the b1 motoneuron inhibited the activity of the b1 muscle, we directly imaged the calcium activity in flight steering muscles using a previously described experimental paradigm (see Materials and Methods)^2^. For these experiments, we co-expressed both i) the genetically encoded calcium indicator GCaMP6f in the wing steering muscles using the driver line *R39E01-LexA* and ii) Kir2.1 in the b1 motoneuron using the *b1-GAL4* driver line. We presented tethered flies with visual stimuli on a LED screen while simultaneously monitoring activity in steering muscles along with wingbeat amplitude and frequency (Fig. S4A). Figure S4B shows examples of pixel-wise variance across entire *∼*200 s bouts of closed-loop stripe fixation for one b1-silenced fly and two genetic control flies. While the genetic control examples show large pixel variance in the b1 muscle area, indicating that these flies modulated activity in the b1 muscle throughout the flight bout, this variance is not present in the b1-silenced flies.

At the population level, we analyzed the wing kinematics and muscle activity across the a set of imaged steering muscles during initiation and cessation of flight during bouts of closed-loop stripe fixation (Fig. S4C). Previous studies have demonstrated that the onset of flight is accompanied by a steep increase in activity across all steering muscles^2^. We observed this increase in activity across all muscles in both genetic control lines; however, as expected, the b1-silenced flies exhibited significantly reduced activity in the b1 muscle during initiation. The lack of b1 activity in b1-silenced flies is also present at flight cessation, when many of the direct steering muscles show a peak in activity before entering quiescence. Interestingly, in Figure S4C b1-silenced flies show an increased level of hg4 activity relative to genetic controls; however, activity in the smaller hg (fourth axillary) muscles are more difficult to de-mix than some of the larger muscles like b1 and b2, so it is unclear if this represents a real effect.

From these experiments, we conclude that chronic silencing of the b1 motoneuron effectively inhibits the activity of the b1 muscle.

### Roll control

In parallel to the pitch perturbation experiments shown in Figure 3, we assayed flies’ response to roll perturbations by applying brief (7 ms) magnetic pulses to flies with ferromagnetic pins glued to their dorsal thorax, oriented normal to their sagittal plane. Experiments were carried out as described in Materials and Methods, with flies glued with pins for pitch and roll released simultaneously into the flight chamber. Figure S3A shows an example of a roll perturbation event, with the 3D model fly in the center showing the measured kinematics and five snapshots in time, and the photomontages from the three high-speed videos projected onto the walls of the illustration.

As described in previous studies^4^, flies control for roll perturbations by modulating the difference in stroke amplitude between their left and right wings to produce counter torques about their roll axis. This left/right stroke amplitude (Δ_LR_Φ) modulation over time (*t*) is well-described by a time-delayed PI controller model (Fig. S3B)^4^, which can be written:

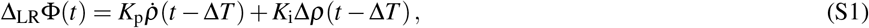

where, as in Equation 1, the controller parameters *K*_p_, *K*_i_, and Δ*T* correspond to the proportional gain, integral gain, and time delay; *ρ* and 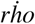 refer to the body roll angle and roll angular velocity, respectively. For each roll perturbation event, we extract the body and wing kinematics from the 3D reconstruction of the video (Materials and Methods) and fit for the control parameters *K*_p_, *K*_i_, and Δ*T*.

Figure S3C shows example roll perturbation events for a genetic control (left) and b1-silenced fly (right). Both flies experience a rightward roll deflection (top) during the period when the magnetic field is applied, indicated by a yellow bar in the time series plots. The bottom row of Figure S3C shows the PI controller fits for these two flies (blue curves) along with the measured difference in left/right stroke amplitude (black dots). At the population level, we compared PI controller model coefficients (*K*_i_, *K*_p_, Δ*T*) between b1-silenced flies and genetic controls for roll control (Fig. S3D). We did not observe any statistically significant differences between the three tested genotypes for any of the control parameters. We also did not observe any significant differences between b2-silenced flies and genetic controls (Fig. S3E,F).

Determining the effects of b1 and b2 silencing on roll control is complicated by the fact that—unlike the case of pitch stabilization— corrective roll maneuvers require an asymmetric response in the left and right wings, while our split-GAL4 driver lines bilaterally silence their targeted motoneurons. However, previous imaging studies have shown that, when presented with fictive roll stimuli, the change in activity of the b1 and b2 muscles is also left/right asymmetric, with activity increasing on the side of increased stroke amplitude, and vice versa^2^. Thus, even with bilateral driver lines, we would expect to see changes in the response to roll perturbations in motoneuron-silenced flies. That we do not see changes to in the roll response indicates that other muscles groups (e.g. the first and third axillaries^2, 14^) likely play a more important role than the basalars in the execution of roll control.

### Quasi-steady aerodynamic model

To estimate the aerodynamic forces and torques produced by the wingbeat patterns observed in our experiments, we used a quasi-steady aerodynamic model developed in previous studies and validated using scaled mechanical models^10, 11, 15^. Specifically, we calculated the total instantaneous quasi-steady aerodynamic forces on a wing, which can be separated into two terms: the translational (**F**_t_) and rotational (**F**_rot_) forces^10, 11, 15, 16^. The total quasi-steady aerodynamic force can thus be written:

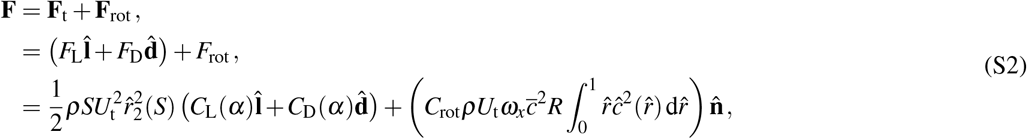

where *ρ* is the air density; *S* the wing area; *U*_t_ the velocity of the wing tip; 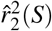 the non-dimensionalized second moment of wing area^17^; *α* is the wing angle of attack; *C*_L_ and *C*_D_ the lift and drag coefficients; *C*_rot_ the rotational force coefficient; *ω*_*x*_ the angular velocity of the wing about its spanwise axis; 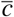 the mean wing chord length; *R* the wing span length; 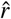 and *ĉ* the non-dimensionalized span and chord; and 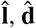, and 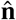 are unit vectors in the directions of the lift, drag, and wing surface normal. We used constant values for the morphological and physical parameters in Equation S2 across flies; the values of these parameters are given in Table S4. The lift and drag coefficients (*C*_L_ and *C*_D_) can be written as functions of the wing angle of attack (*α*)^18^:

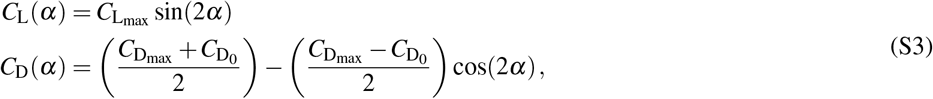

where 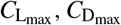, and 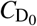 are dimensionless constants best fit for *Drosophila* flight (see Table S4).

To calculate the aerodynamic torque, **T**, produced by a flapping wing, we use the approximation that the wing center of pressure is located 70% along the length of the wing span^19^, and write:

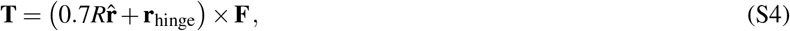

where, as above, *R* is the wing span length, 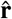 is the wing span unit vector, **F** is the total aerodynamic force produced by the wing, and **r**_hinge_ is the vector from the fly center of mass to the wing hinge (see Table S4). Because Equation S4 is written in the body-tied frame of reference, the pitching torque, *T*_pitch_, corresponds to the *y* component of *T*. As we are primarily interested in the pitch torque, *T*_pitch_, we write the hinge vector, **r**_hinge_, as purely along the fly’s long body axis 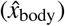, i.e. 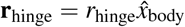. When presenting aerodynamic force calculations (Fig. S2), we normalize these values to the weight of the fly, i.e. **F** *→* **F***/mg*. Similarly, we normalize aerodynamic torque by the product of the fly’s weight and wing span length, i.e. **F** *→* **T***/mgR* (Fig. 2,S2).

### Flight simulation

To simulate flapping flight, we used a framework similar to the one described in^5^. In these simulations, we numerically solve the Newton-Euler equations of motion in the longitudinal plane, i.e. we restrict a simulated fly’s motion to two translational and one rotational degrees of freedom, so that the state vector describing the fly’s body can be written:

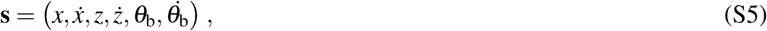

where *x* and *z* denote the body center of mass coordinates in the forward and vertical directions, and *θ*_b_ the body rotation angle about the pitch axis. The dot notation (e.g. 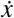) refers to the derivative with respect to time of the given variable. In these coordinates, we write the Newton-Euler equations for rigid body motion in the fly body frame:

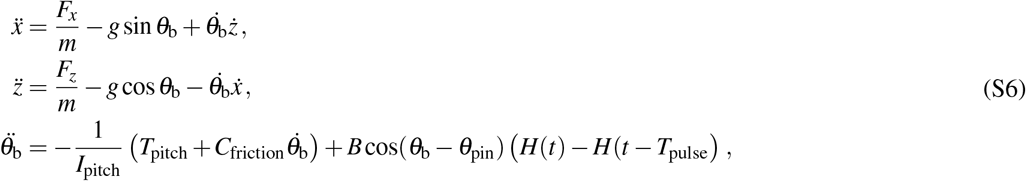

where *m* is the fly body mass; *I*_pitch_ is the pitch moment of inertia; *g* is the gravitational acceleration; *F*_*x*_ and *F*_*z*_ are the *x* and *z* components of aerodynamic forces generated by the flapping wings; *T*_pitch_ is the component of aerodynamic torque generated by the wings; and *C*_friction_ is the coefficient of pitch rotational drag. The last term on the right hand side of the equation for 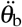 is the simulation equivalent of the magnetic perturbation applied in our experiments (Fig. 3): *H* represents the Heaviside step function, so this term is nonzero only during the duration of the applied perturbation, *t* ∈ [0, *T*_pulse_]. The angle that the magnetic pin makes with the fly body in the sagittal plane, *θ*_pin_, is kept fixed throughout, while the magnetic perturbation strength, *B*, is kept constant for a given simulation run, but varied when the simulation is fit to data. Table S4 gives a list of the constant parameters used in our simulations.

To calculate the aerodynamic forces and torques generated by the flapping wings, we prescribed the wing kinematics of the simulated fly using a set of parameterized equations, as in^5, 9^, and used a quasi-steady aerodynamic model to determine the forces generated by these flapping patterns (see Eq. S2 above). The parameterized form for the wing kinematics is given by:

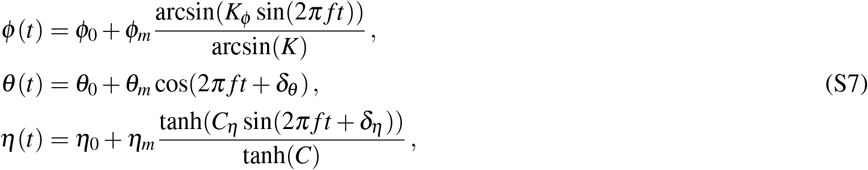

where *ϕ* (*t*), *θ* (*t*), and *η*(*t*) are the wing Euler angles—stroke, deviation, and rotation (see Fig. 1F)—as a function of time; *ϕ*_0_, *θ*_0_, and *η*_0_ are angle offsets; *ϕ*_*m*_, *θ*_*m*_, and *η*_*m*_ are amplitudes; *δ*_*θ*_ and *δ*_*η*_ are phase offsets; *K*_*ϕ*_ and *C*_*η*_ are dimensionless parameters that tune the shape of the waveform; and *f* is the wingbeat frequency. For our flight simulations, all parameters but *ϕ*_0_ and *ϕ*_*m*_, which are used to set the forward stroke amplitude, were kept fixed and based on an optimization for hovering stability within our simulation framework, with the deviation angle, *θ*, set to 0° for simplicity (see Table S4). For a given set of wing Euler angles, we write the unit vectors pointing in the direction of the wing span and chord (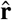 and **ĉ**, respectively)—used to determine wingtip velocity, torque lever arm, etc. in Equations S2,S4—as:

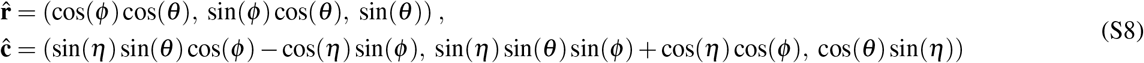

The simulated fly implements PI control for its pitch degree of freedom as in Equation 1, which computes the change in forward stroke angle, Δ_fwd_*ϕ*. In terms of the wing kinematic parameters in S7, this corresponds to an added change to the stroke angle parameters *ϕ*_0_ and *ϕ*_*m*_ given by:

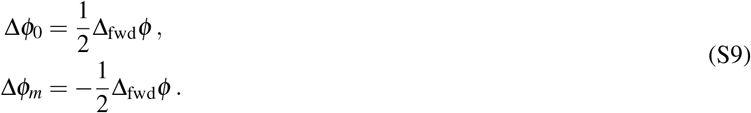

Because Equation 1 is defined for continuous time, we modify the time delay so that the value of Δ_fwd_*ϕ* is updated only once per wingstroke.

Using this simulation framework, we performed fits to experimental data by allowing the controller parameters *K*_i_, *K*_p_, and Δ*T*, as well as the magnetic perturbation strength, *B*, to vary and perform a least squares fit to measured body kinematics (see Materials and Methods).

### Simplified closed-loop model

Mechanical dynamics are crucial to locomotor control^20–22^, so we further assessed the closed-loop dynamics of *Drosophila* pitch control in the presence of motoneuron manipulations by estimating a linearized plant model, and using the result to examine the closed-loop dynamics. Beginning with the equation of motion for the pitch degree of freedom (Eq. S6), we consider the unforced (*B* = 0) dynamics. We approximate the torque, *T*_pitch_, as a linear combination of the delayed PI feedback Δ_fwd_*ϕ* (Eq. 1) and body pitch angle. This gives:

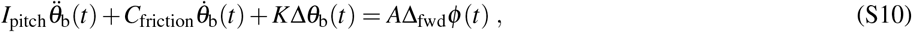

where Δ_fwd_*ϕ* (*t*) is given by the PI controller in Eq. 1. The first fitting parameter, *K*, is analogous to a spring constant and captures the relationship between body pitch and torque that can arise, for example, due to an offset between the center of lift and center of mass^23, 24^. The second fitting parameter, *A*, captures the scaling that relates changes in wing kinematic output to changes in pitch torque. The remaining terms in Equation S10 are defined as above: *I*_pitch_ and *C*_friction_ are the moment of inertia and coefficient of rotational drag about the pitch axis, respectively (Tab. S4); and 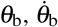, and 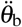 are the body pitch angle, velocity, and acceleration.

Using Equation S10, measured body pitch kinematics, fitted controller model parameters, and literature values for *I*_pitch_ and *C*_friction_^25^, we performed a least-squares fit for the spring constant, *K*, and scaling term, *A*. With the resultant fit parameters (*K*=7.25 *×* 10^*−*10^ Nm, *A*= *−*8.28 *×* 10^*−*9^ Nm), the plant and controller transfer functions in our linearized system—*P* and *C*, respectively—can be written:

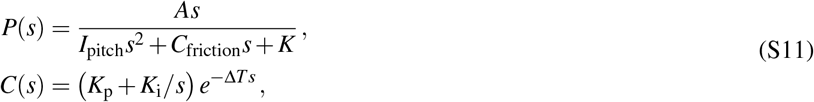

where *s* is the Laplace transform complex frequency and *e*^*−*Δ*Ts*^ represents the controller time delay. Here, *P*(*s*) is taken to be the transfer function from Δ_fwd_*ϕ* (*t*) to 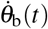. Using MATLAB’s `feedback` command, we generated the closed-loop transfer function *G*(*s*) = *P*(*s*)*/*(1 *− P*(*s*)*C*(*s*)), eliminating the pole–zero cancellation at *s* = 0 using MATLAB’s `minreal`; *G*(*s*) maps torque perturbations to pitch angular velocities. We used `lsim` with *G* to simulate responses to a simulated 15 ms magnetic pulse, and integrated the result using MATLAB’s `cumtrapz` command to calculate Δ*θ*_b_(*t*). We tuned the impulse magnitude, i.e. the magnetic field strength *B*, by hand until the output of the linearized dynamics had approximately the same amplitude as the data and nonlinear simulation. We found that the resulting value of *B* matched the range of values obtained for *B* in our nonlinear fitting.^1^ The time course of simulated linearized dynamics showed excellent qualitative agreement with both our behavioral data and flapping flight simulations (Fig. S9A).

The accuracy with which this linearized model captured our experimentally observed dynamics allowed us to further explore the effects of differing control strategies on stability. In particular, using Equation S10, we calculated the damping ratio, *ζ*, for each fly genotype (Fig. S9B), under a simplifying assumption of no time delay in the feedback loop. As shown in Figure S9B, the closed-loop dynamics for genetic control flies were slightly underdamped (*ζ ≈* 0.63). For b1-silenced flies, the dynamics were slightly overdamped (*ζ >* 1), resulting in a slower system response. For b2-silenced flies, the system became sufficiently underdamped to go unstable when the time delay was reintroduced, although only marginally so, resulting in a very slow unstable mode. Estimating open-loop transfer functions can sensitively depend on the closed-loop dynamics^26^, and we suspect that this instability may be an artifact of the relatively short time window over which we measured kinematic data, thus fitting short-term transients while not capturing steady-state stability (we made no effort to constrain the parameter estimates to ensure stability). This linear analysis highlights the critical role of velocity feedback^27^ (i.e. analogous to a traditional “derivative controller,” since the mechanosensors produce delayed angular-velocity-dependent feedback^28, 29^) on pitch stability in addition to the importance of integral control of pitch velocity (i.e. “proportional” to change in body-pitch angle) on transient performance.

### Motoneuron driver lines

Because the *b1-GAL4* (*MB258C-GAL4*) driver line contained off-target VNC expression in the neck tectulum and abdominal neuromere^30^, we performed optogenetic excitation and inhibition experiments as in Figure 2 using the split-GAL4 line *SS04528-GAL4*, which targets also targets the b1 motoneuron but does not share common off-target expression with *b1-GAL4*^31^. Figure S5 shows the expression pattern of *SS04528-GAL4* in the brain and VNC (c.f. Fig. S1A,B), as well as the body and wing kinematic changes evoked via optogenetic excitation (Fig. S5C–E) and silencing (Fig. S5F–G) compared to both *b1-GAL4* and the empty split *SS01062-GAL4*. The body and wing responses to both excitation and silencing between *SS04528-GAL4* and *b1-GAL4*—light and dark blue in Figure S5, respectively—are in agreement. We observed the same phenotypic similarity when comparing the full set of wing Euler angles for wingbeats prior to and during the LED stimulus (Fig. S5). Taken together, these results indicate that the flight phenotypes we observed using *b1-GAL4* are indeed due to manipulation of the b1 motoneuron, and not off-target cells. We were unable to find an alternate driver line targeting the b2 motoneuron to perform a complementary tests of *b2-GAL4*; however, *b2-GAL4*’s relatively low level of off-target expression (Fig. 1C, S1C,D) and the agreement between our results (Fig. 2, S2) and previous electrophysiology/imaging studies^2, 32, 33^ suggest that the phenotypes we observed are very likely due to b2 motoneuron manipulation.

### Combined optogenetic and mechanical perturbation

As an alternative to chronic silencing, we performed experiments in which we crossed our motoneuron driver lines to *UAS-GtACR1* in order to perform combined optogenetic silencing and magnetic perturbations. We performed two versions of these experiments. In the first, both LED and magnetic field were applied simultaneously for 15 ms, as in the case of the data used to fit for simulation parameters in Figure 3H,I. In the second version, the two signals differed in both their duration and onset time—specifically, we used a 50 ms LED pulse for optogenetic silencing followed by a 7 ms magnetic field pulse that began 15 ms after the onset of the LED. For this latter paradigm, we performed fit PI controller models to the pitch and roll stabilization responses (Fig. S7), as in the analysis of chronic silencing data. Again, our findings using this combined optogenetic and magnetic perturbation were consistent with the results of the chronic silencing experiments, only with a smaller effect size: b1- and b2-silenced flies showed decreases integral and proportional gains for pitch control, respectively (Fig. S7A), and we observed no effect on roll control (Fig. S7B).

## Supplementary Figures

**Figure S1.**
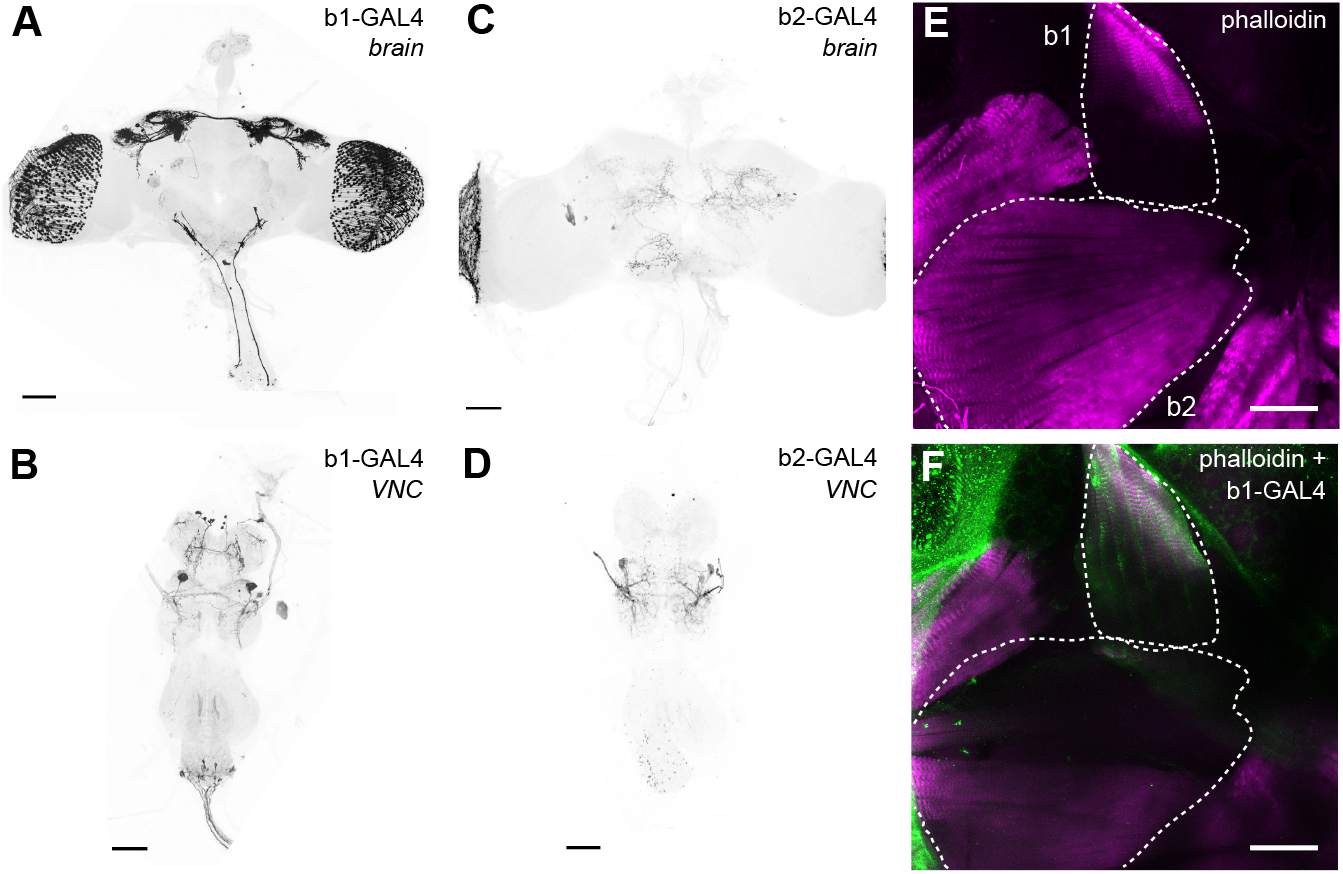
Maximum intensity projection (MIP) images of the split driver lines *b1-GAL4* and *b2-GAL4*. **A** Maximum intensity projection (MIP) brain image from a *b1-GAL4 > CsChrimson* fly. Black corresponds to mVenus, light gray to DNCad (neuropil). **B** MIP VNC image from a *b1-GAL4 > CsChrimson* fly. Same image as in Figure 1B. **C**,**D** Same as **A**,**B** but with *b2-GAL4 > CsChrimson* flies. **D** shows the same image as in Figure 1C. **E**,**F** Phalloidin-stained thoracic hemisection from a *UAS-GFP > b1-GAL4* fly showing wing musculature. b1 and b2 muscles outlined with white dashed lines. **E** shows just the phalloidin channel (magenta) across the full stack; **F** shows the MIP of a partial stack with both phalloidin and GFP (green) expression innervating the b1 muscle. All scale bars 50 μm. See Table S3 for full fly genotypes.

**Figure S2.**
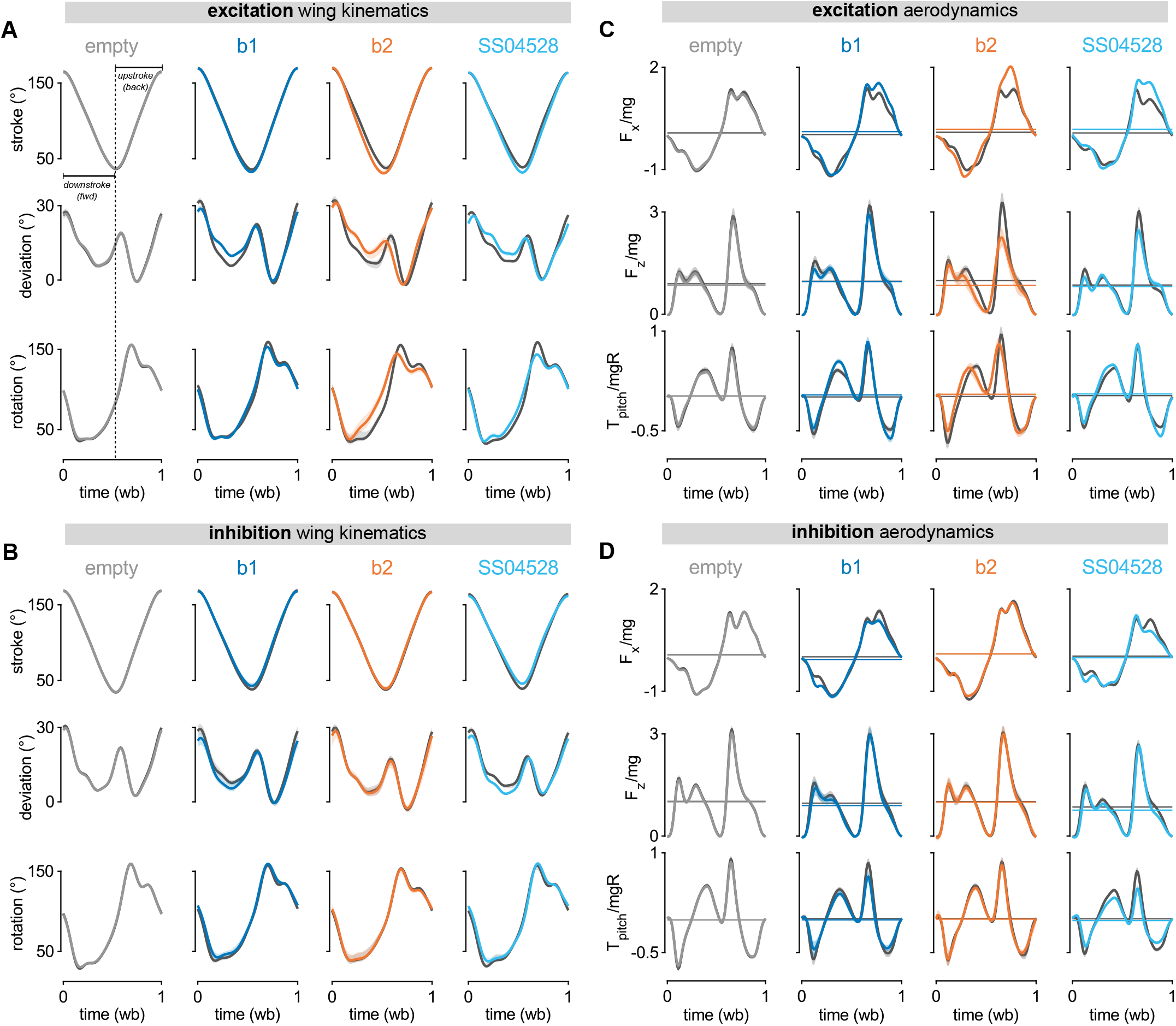
Extended results on the optogenetic activation and silencing of the b1 and b2 motoneurons. **A** Population-averaged time courses of the three wing Euler angles (stroke, deviation, and rotation) comparing pre- (dark gray) and during-LED-stimulus (light gray, blue, orange) across a single wingbeat for bouts of optogenetic excitation with CsChrimson. Columns from left to right: *empty* (aka *SS01062-GAL4*; light gray), *b1-GAL4* (dark blue), *b2-GAL4* (orange), and *SS04528-GAL4* (light blue/cyan; targets b1 motoneuron). **B** Same as in **A** but for bouts of optogenetic silencing with GtACR1. **C** Population-averaged time courses of normalized quasi-steady aerodynamic forces and torques comparing pre-(dark gray) and during-LED-stimulus (light gray, dark blue, orange, light blue/cyan) across a single wingbeat for bouts of optogenetic excitation with CsChrimson. Rows from top to bottom: forward/backward force (*F*_*x*_*/mg*), up/down force (*F*_*z*_*/mg*), and pitch torque (*T*_pitch_*/mgR*). Columns correspond to the empty, b1, and b2 genotypes as in **A**,**B**. Horizontal lines show wingbeat-averaged values of force or torque. **D** Same as **C** but for bouts of optogenetic silencing with GtACR1. In all plots, times series curves and envelope correspond to population mean *±*95% confidence interval (500-sample bootstrap). See Table S3 for full fly genotypes.

**Figure S3.**
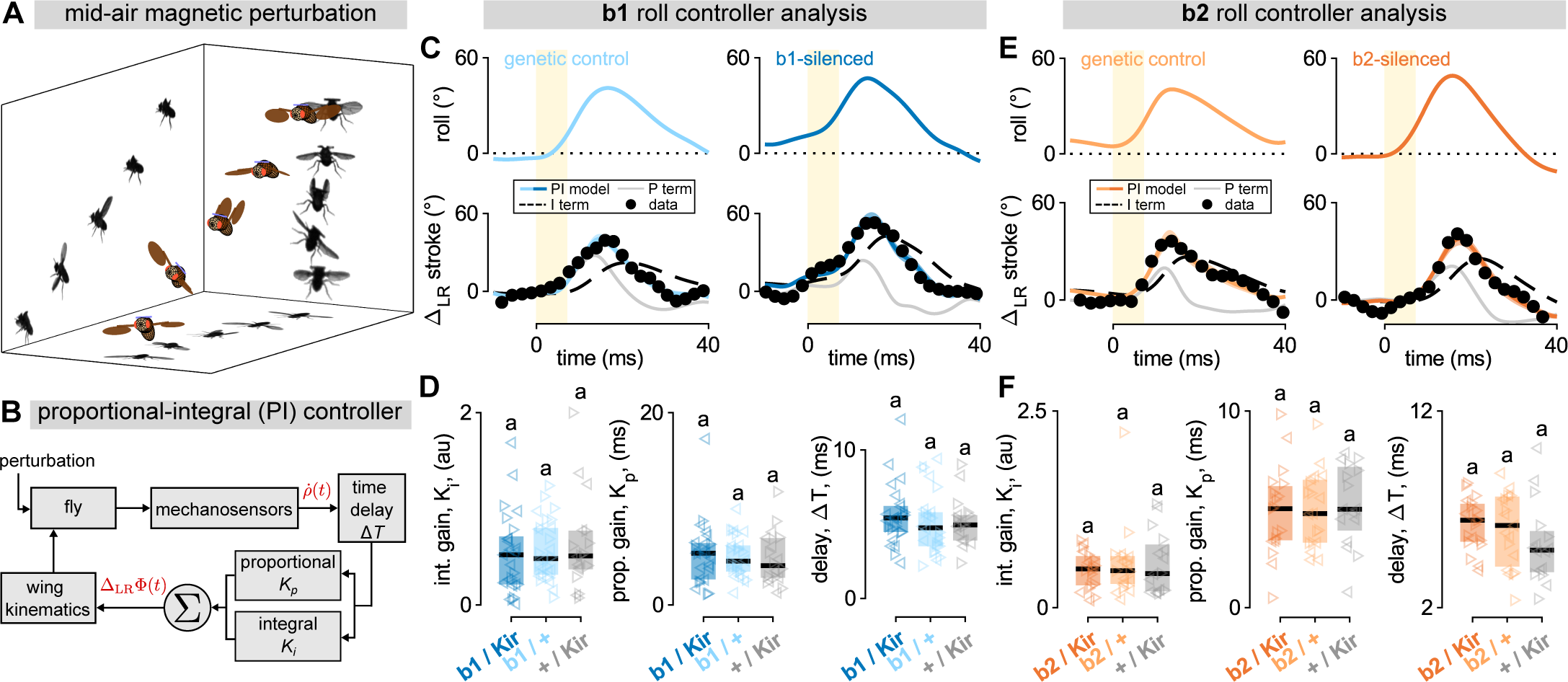
Chronic b1 and b2 inhibition do not alter roll stabilization behavior. **A** Illustration of a fly experiencing and correcting for a leftward roll perturbation. The 3D rendered fly represents measured kinematics, with attached ferromagnetic pin false-colored blue. Walls show photomontages from the three high-speed cameras filming the perturbation event. **B** Generalized proportional-integral (PI) controller model for rapid flight stabilization, as in Equation S1 and Figure 3B. **C** Example roll right perturbation analyses for a genetic control fly (left) and a b1-silenced fly (right). Top row shows change in body roll angle over time (blue traces); bottom row shows measured change in left-right stroke amplitude difference over time (black dots) as well as PI controller model fit to data (blue traces with 95% CI). Bottom row plots also show contributions of individual model terms: P term (proportional; thin gray line) and I term (integral; dashed black line). Yellow bar indicates 7 ms magnetic pulse. **D** Summary statistics for PI controller model parameters from Equation S1—integral gain (*K*_i_; left), proportional gain (*K*_p_; center), and time delay (Δ*T*, right)—for b1-silenced flies (dark blue; *N*=25) and two genetic controls (light blue and gray; *N*=26 and 18). Each data point represents a roll perturbation movie, with left- and right-pointing triangles representing left and right roll perturbations, respectively. Horizontal black lines and boxes show median and interquartile range for each genotype. **E** Same as in C but with a genetic control (left) and b2-silenced fly (right). **F** Same as **D** but with b2-silenced flies (dark orange; *N*=19) and two genetic controls (light orange and gray; *N*=19 and 16). Lower case letters above data in both **D** and **F** indicate significance categories, determined via Wilcoxon rank sum test with Bonferroni correction (*α*=0.05). See Table S3 for full fly genotypes.

**Figure S4.**
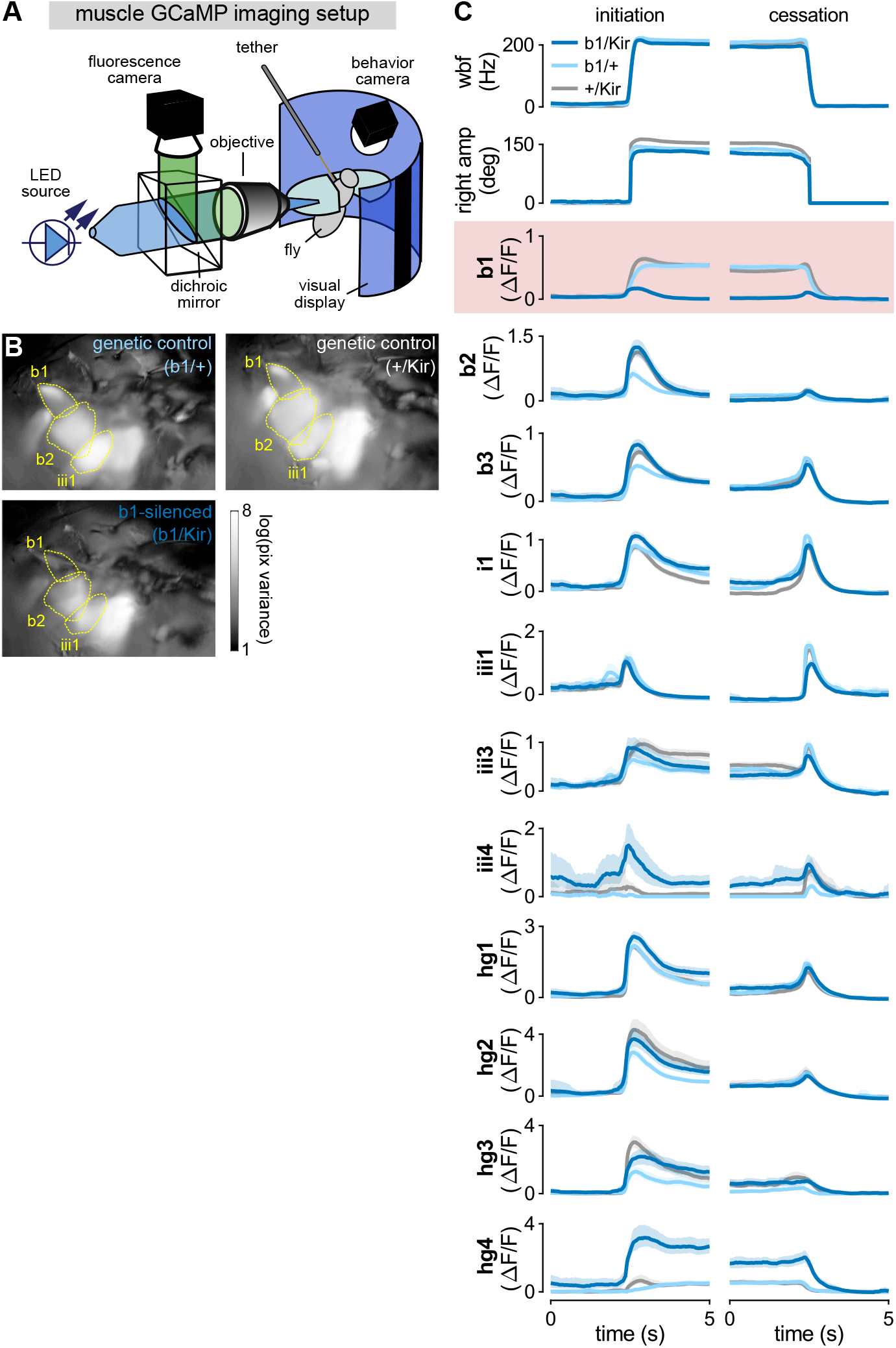
Calcium imaging of muscle activity. **A** Schematic for muscle imaging apparatus. The steering muscles on the right side of tethered flies are imaged using epifluorescent optics while the flies are presented with visual stimuli on the display screen. Simultaneously the stroke amplitude and wingbeat frequency are also recorded. Schematic from^2^. **B** Pixel variance in steering muscle imaging across ∼200 seconds of continuous closed loop stripe fixation for genetic controls (top row) and b1-silenced flies (bottom). Yellow dashed lines show outlines of the b1, b2, and iii1 muscles as reference. **C** Mean wing kinematics and muscle activity at the initiation (left) and cessation (right) of flight for b1-silenced flies (dark blue; *N*=18 flies) and two genetic controls (light blue and gray; *N*=17 and *N*=20 flies). See Table S3 for full fly genotypes.

**Figure S5.**
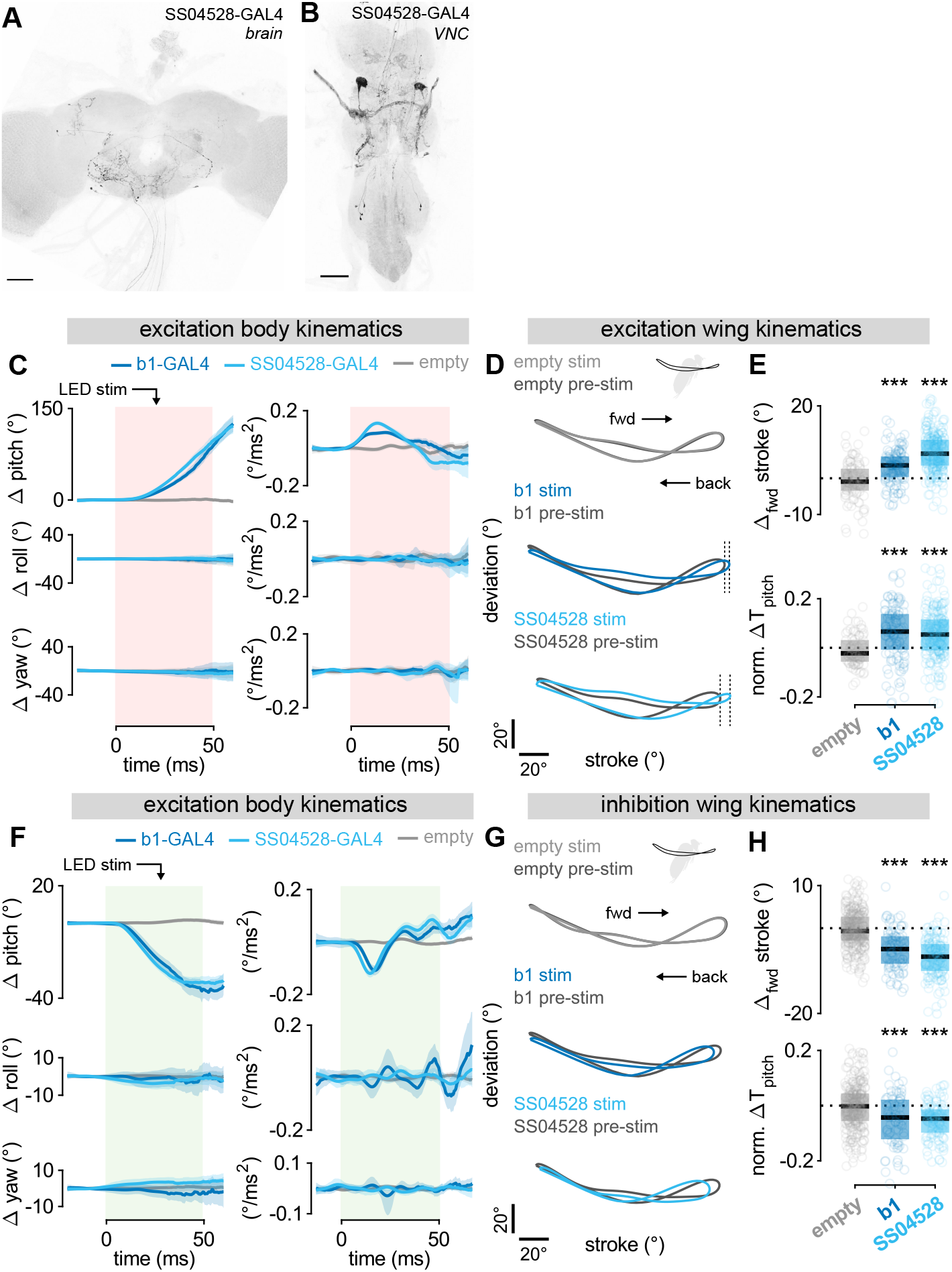
Two driver lines targeting the b1 motoneuron respond consistently to optogenetic manipulation. **A** Maximum intensity projection (MIP) image of the brain of a *UAS-CsChrimson > SS04528-GAL4* fly. Black corresponds to mVenus, light gray to DNCad (neuropil). Scale bar 50 μm. **B** MIP image of a *UAS-CsChrimson > SS04528-GAL4* fly VNC. Colors and scale bar as in **A. C** Body kinematics versus time in response to 50 ms optogenetic activation of *b1-GAL4* (dark blue; *N*=140 movies), *SS04528-GAL4* (light blue/cyan; *N*=272 movies), and *SS01062-GAL4* (aka *empty*; gray; *N*=108 movies) flies with *UAS-CsChrimson*, as in Figure 2A. Rows correspond to rotational degrees of freedom: pitch (top), roll (middle), and yaw (bottom). Columns give angular displacement (left) and angular acceleration (right). Data shown represent mean *±*95% confidence interval. **D** Wing kinematic data for movies in **C**. Plots show wing tip angular position in the wingstrokes prior to (dark gray; “pre-stim”) and during (light gray, dark blue, light blue/cyan; “stim”) optogenetic activation. Thick traces represent population averages; thin lines represent single-fly wingbeats. Vertical and horizontal scale bars provide 20° references for deviation and stroke angles, respectively. **E** Change in forward stroke angle (top) and normalized, wingbeat-averaged aerodynamic pitch torque (bottom) for wingbeats prior to and during optogenetic activation of *b1-GAL4* (dark blue), *SS04528-GAL4* (light blue/cyan), and *SS01062-GAL4* (gray) flies. Circles show raw data; box and horizontal line show interquartile range and median, respectively. Statistical significance determined via Wilcoxon signed-rank test (***, *p*<0.001; **, *p*<0.01; *, *p*<0.05). **F**–**H** Same as in **C**–**E** but with *b1-GAL4* (dark blue; *N*=89 movies), *SS04528-GAL4* (light blue/cyan; *N*=146 movies), and *SS01062-GAL4* (gray; *N*=323 movies) flies. See Table S3 for full fly genotypes.

**Figure S6.**
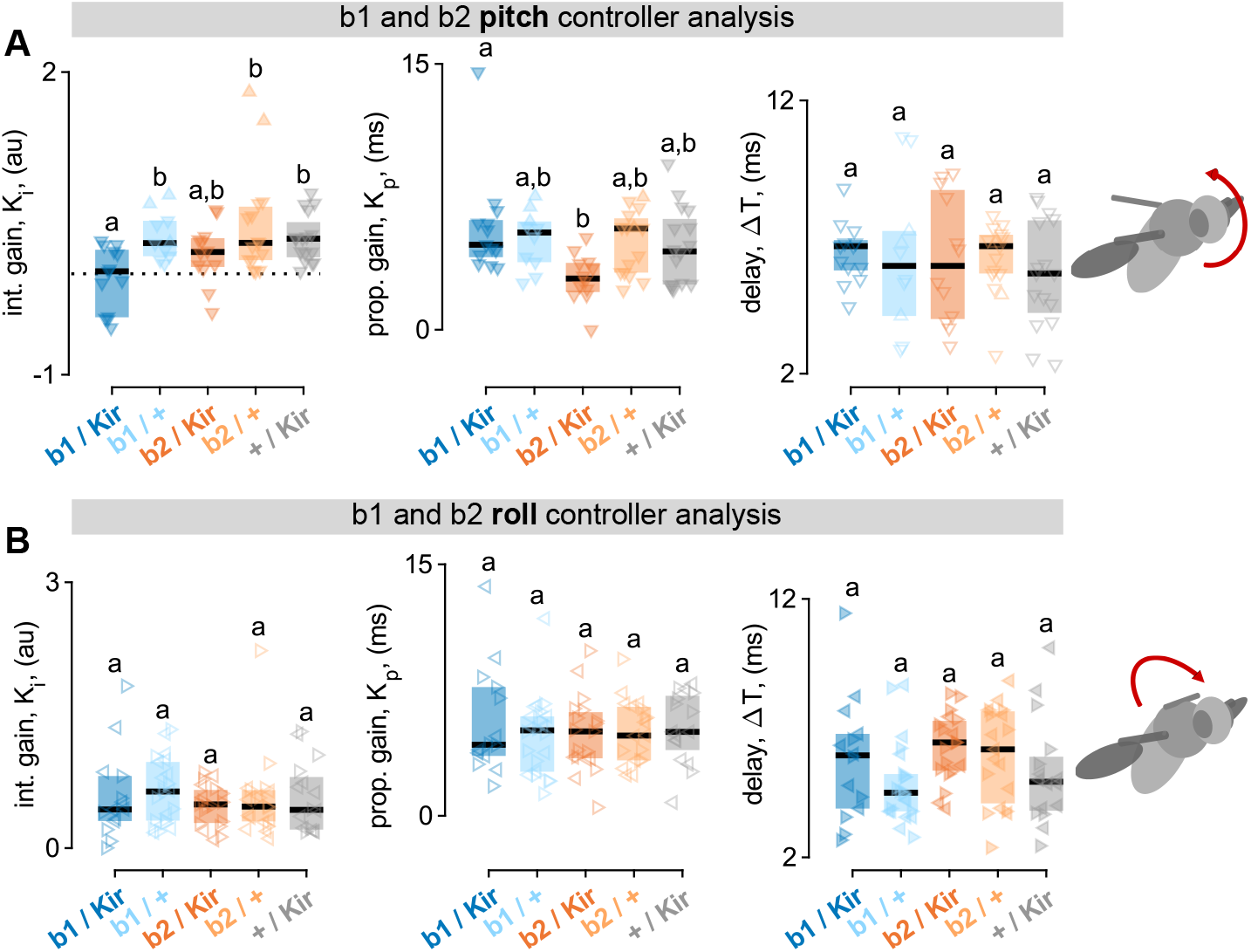
Perturbation experiments with *5XUAS-Kir2*.*1* effector line. **A** PI controller coefficients—*K*_i_(integral gain, left), *K*_p_(proportional gain, middle), Δ*T* (time delay, right)—for fits to pitch perturbation movies as in Figure 3D,F. Data shown for *b1/Kir2*.*1* (dark blue; *N*=18 movies), *b1/+* (light blue; *N*=16 movies), *b2/Kir2*.*1* (dark orange; *N*=17 movies), *b2/+* (light orange; *N*=22 movies), and *+/Kir2*.*1* (gray; *N*=22 movies). Data for b2 flies and genetic controls is the same as in Figure 3F. Lower case letters above data indicate significance categories, determined via Wilcoxon rank sum test with Bonferroni correction (*α*=0.05). **B** Same as in **A** but for roll correction videos. Data shown for *b1/Kir2*.*1* (dark blue; *N*=14 movies), *b1/+* (light blue; *N*=22 movies), *b2/Kir2*.*1* (dark orange; *N*=19 movies), *b2/+* (light orange; *N*=19 movies), and *+/Kir2*.*1* (gray; *N*=16 movies). Data for b2 flies and genetic controls is the same as in Figure S3F. See Table S3 for full fly genotypes.

**Figure S7.**
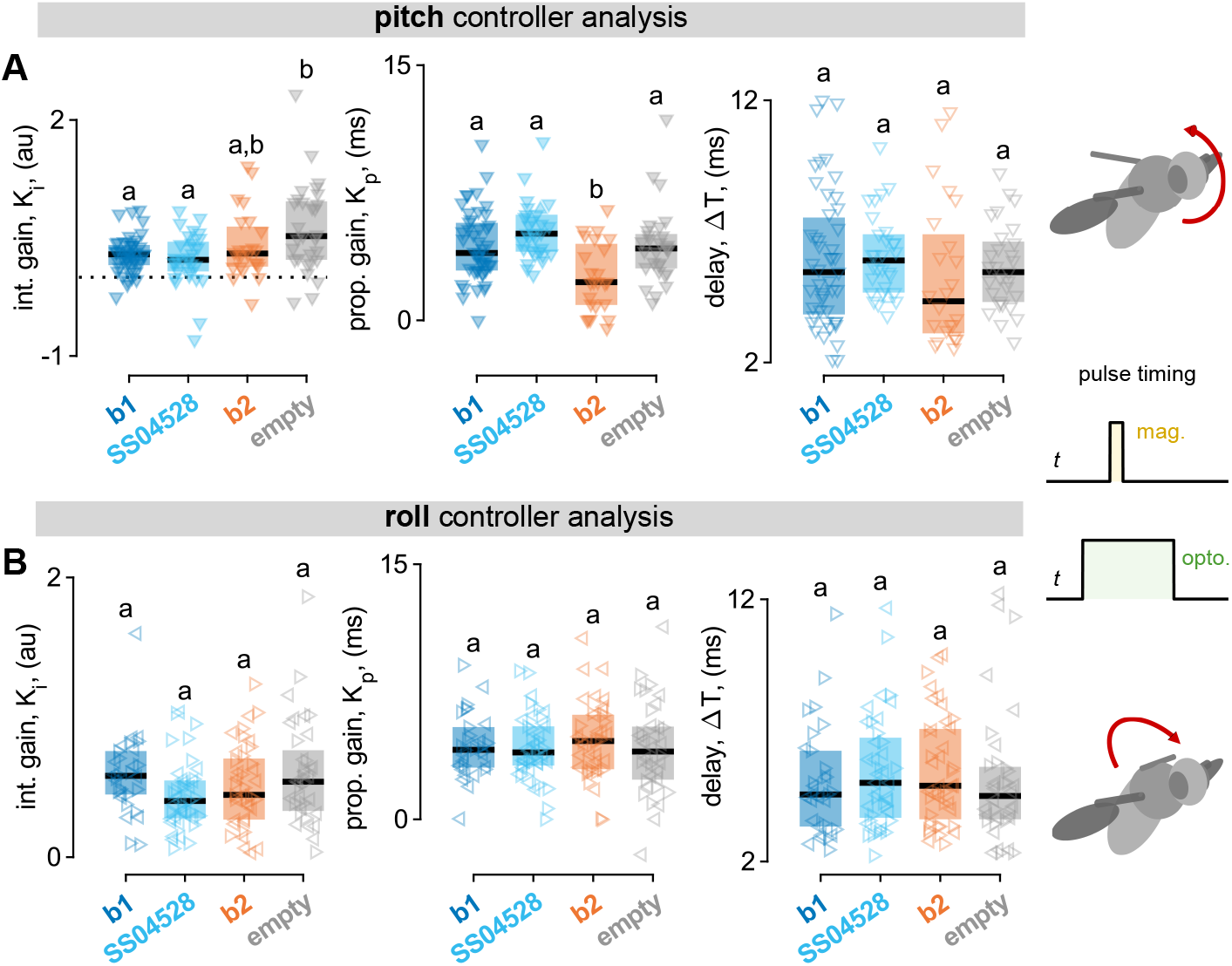
Combined optogenetic silencing and mechanical perturbation experiments. **A** PI controller coefficients—*K*_i_(integral gain, left), *K*_p_(proportional gain, middle), Δ*T* (time delay, right)—for fits to pitch perturbation movies wherein flies are exposed to a 50 ms LED pulse to induce optogenetic silencing followed by a 7 ms magnetic perturbation that begins 15 ms after the LED onset (pulse structure schematized on the right). Data shown for *UAS-GtACR1* crossed with *b1-GAL4* (dark blue; *N*=47 movies), *SS04528-GAL4* (alternate b1 MN driver line; light blue/cyan; *N*=30 movies), *b2-GAL4* (orange; *N*=24 movies), and *SS01062-GAL4* (aka *empty*; gray; *N*=29 movies) flies. Lower case letters above data indicate significance categories, determined via Wilcoxon rank sum test with Bonferroni correction (*α*=0.05). **B** Same as in **A** but for roll correction videos with the same structure of optogenetic pulse and magnetic perturbation. Data shown for *b1-GAL4* (dark blue; *N*=30 movies), *SS04528-GAL4* (alternate b1 MN driver line; light blue/cyan; *N*=40 movies), *b2-GAL4* (orange; *N*=40 movies), and *SS01062-GAL4* (aka *empty*; gray; *N*=36 movies) flies. See Table S3 for full fly genotypes.

**Figure S8.**
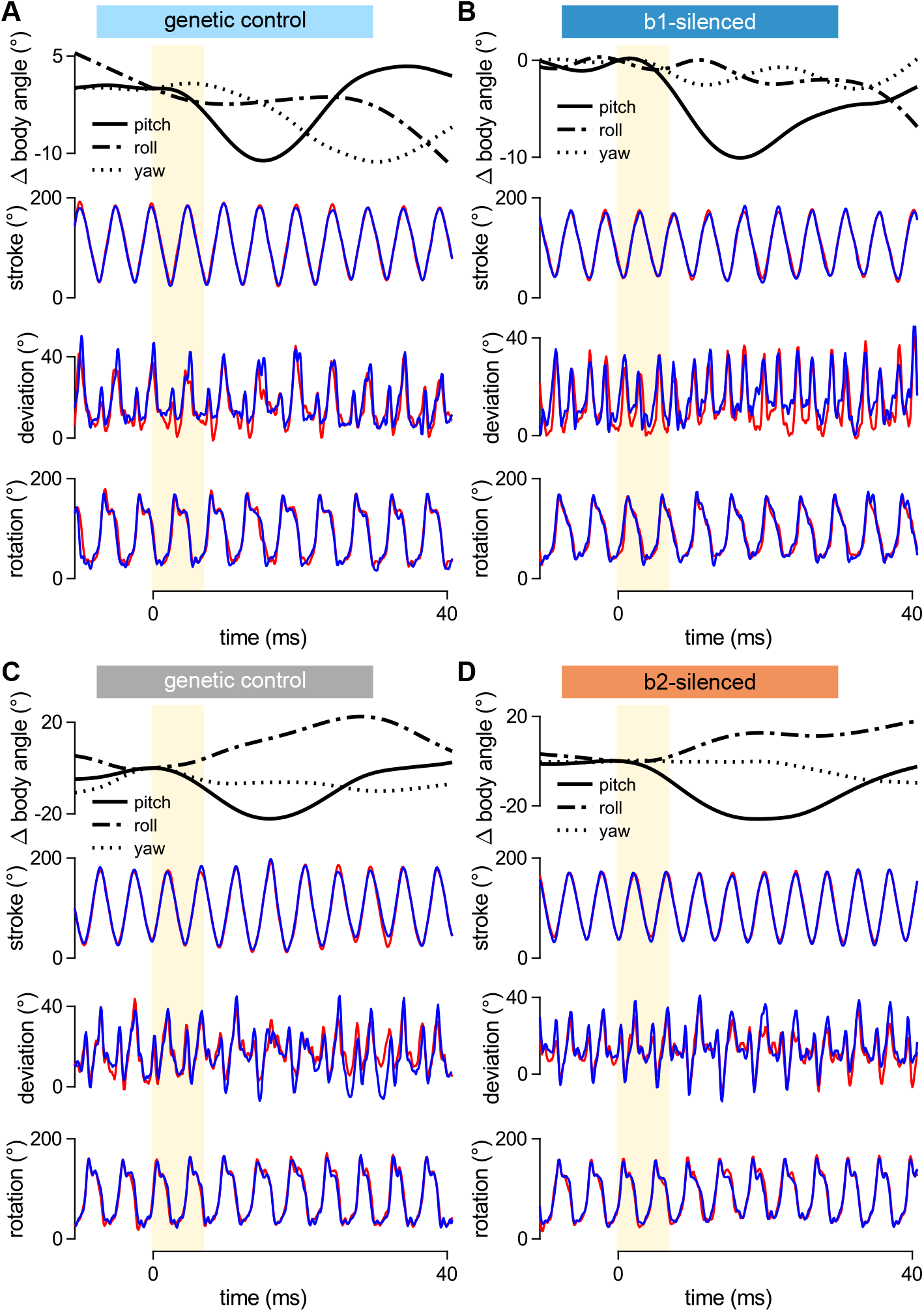
Full kinematics for example perturbations. Related to Figure 3. **A**–**D** Body and wing Euler angles as a function of time for example perturbation events shown in Figure 3C,E. In each panel, top row show the body Euler angles pitch (solid lines), roll (dashed-dotted lines), and yaw (dotted lines). Bottom three rows show wing Euler angles (stroke, top; deviation, middle; rotation, bottom), with red and blue lines denoting right and left wing, respectively. Yellow bar indicates period of 7 ms magnetic pulse. See Table S3 for full fly genotypes.

**Figure S9.**
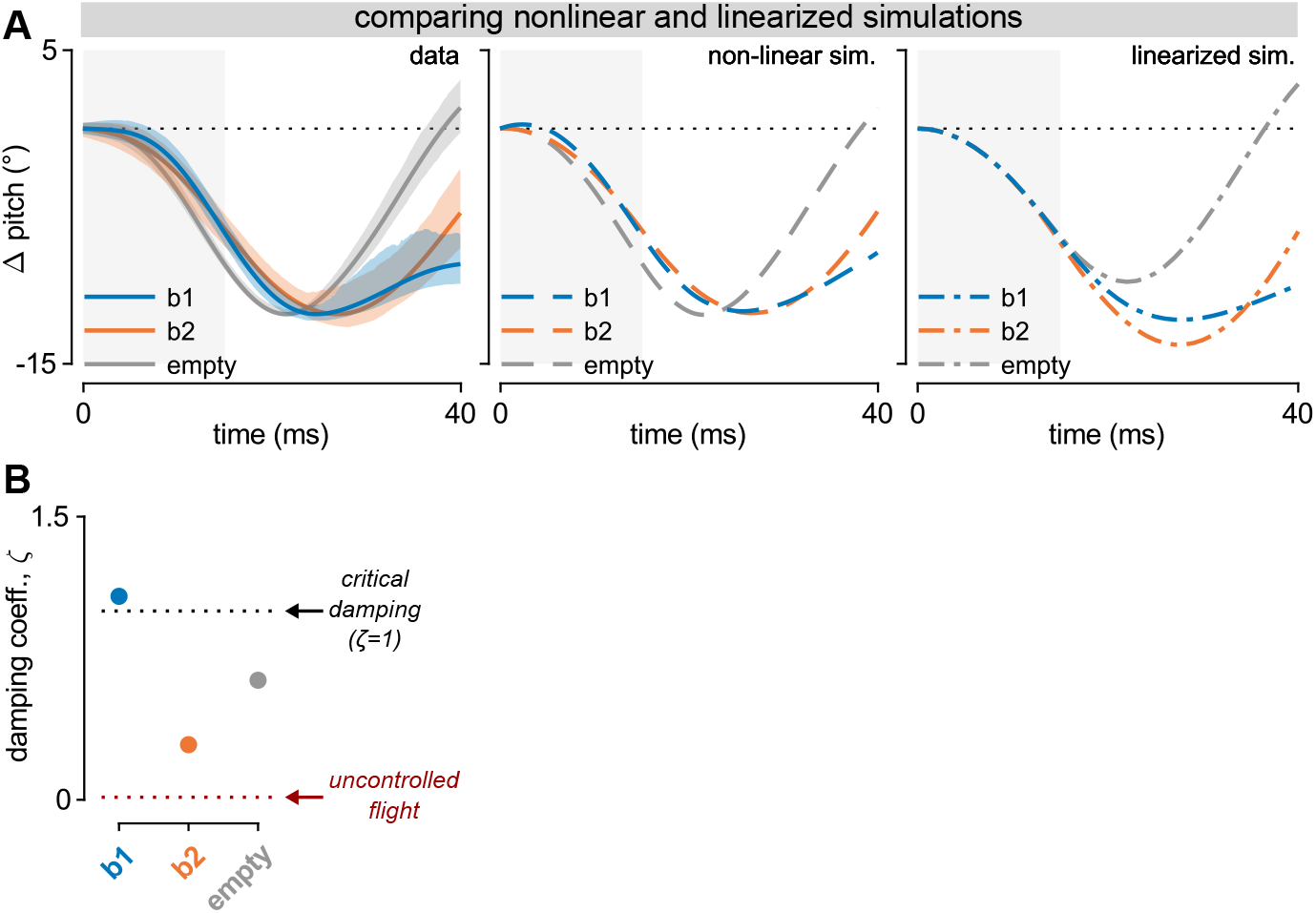
Linearized model for closed-loop dynamics. **A** Change in body pitch angle over time as measured from experimental data (population average with 95% CI envelope; left), flapping flight simulation (middle), and linearized dynamical model (right). Curve colors correspond to different genotypes: *b1-GAL4 > GtACR1* (“b1”; blue), *b2-GAL4 > GtACR1* (“b2”; orange), and *empty > GtACR1* (“empty”; gray). Gray bar represents the 15 ms simultaneous LED and magnetic field stimuli, which optogenetically silence and impose external torque, respectively. Left and middle plots are the same data as in Figure 3H. **B** Damping ratio, *ζ*, estimated from linearized model of closed-loop dynamics for b1-silenced (blue), b2-silenced (orange), and genetic control (gray) flies. See Table S3 for full fly genotypes.

## Supplementary Tables

**Table S1.** Fly stocks.

| Name | Genotype | Source |
| --- | --- | --- |
| *b1-GAL4 (MB258C)* | R71D08-p65ADZp (attP2), R33E02-ZpGAL4DBD (VK00027) | gift from Troy Shirangi |
| *b2-GAL4* | VT40232-p65ADZp (attP40) ; VT23830-ZpGAL4DBD (attP2) | gift from Anne von Phillipsborn<sup>34</sup> |
| *empty (SS01062)* | R24A03-p65ADZp (attP40) ; R74C01-ZpGdbd (attP2) | gift from Gwyneth Card<sup>31</sup> |
| *SS04528* | R19G08-p65ADZp (attP40) ; R47F01-ZpGDBD (attP2) | gift from Gwyneth Card<sup>31</sup> |
| *UAS-CsChrimson* | 20XUAS-CsChrimson-mVenus (attP18) | gift from Gwyneth Card<sup>35</sup> |
| *UAS-GtACR1* | pJFRC7-20XUAS-IVS-GtACR1-EYFP (attP40) | gift from Nilay Yapici<sup>36</sup> |
| *10XUAS-Kir2.1* | pJFRC49-10XUAS-IVS-eGFPKir2.1 (attP2) | gift from Troy Shirangi |
| *5XUAS-Kir2.1* | pJFRC49-5XUAS-IVS-eGFPKir2.1 (attP2) | gift from David Stern |
| *UAS-GFP* | 10XUAS-IVS-mCD8::GFP (attP40) | BDSC #32186 |
| *R39E01-LexA* | R39E01-LexA (attP40) | BDSC #52776 |
| *LexAop-GCaMP6f* | 13XLexAop2-IVS-GCaMP6f-p10 (su(Hw)attP5) | BDSC #44277 |
| *b1-GAL4 ∩ tshLexA* | tshLexA (attP40) ; R71D08-p65ADZp (attP2), R33E02-ZpGAL4DBD (VK00027) | from Troy Shirangi |
| *8XLexAop2-FlpL ∩ 10XUAS-FRT>STOP>FRT-Kir2.1* | w+ ; pJFRC79-8XLexAop2-FlpL (attP40) ; pJFRC56-10XUAS-FRT > STOP > FRT-kir2.1-gfp (attP2) | from Troy Shirangi |

**Table S2.** Full genotype of flies used in experiments.

| Data shown in | Experiment type | Abbreviated genotype | Full genotype |
| --- | --- | --- | --- |
| Figure 1B | VNC anatomy | *b1-GAL4* | *20XUAS-CsChrimson-mVenus (attP18)/+ ; +/+ ; R71D08-p65ADZp (attP2) , R33E02-ZpGAL4DBD (VK00027)/+* |
| Figure 1C | VNC anatomy | *b2-GAL4* | *20XUAS-CsChrimson-mVenus (attP18)/+ ; VT40232-p65ADZp (attP40)/+ ; VT23830-ZpGAL4DBD (attP2)/+* |
| Figure 1G | opto. activation | *b1-GAL4 > CsChrimson* | *20XUAS-CsChrimson-mVenus (attP18)/+ ; +/+ ; R71D08-p65ADZp (attP2) , R33E02-ZpGAL4DBD (VK00027)/+* |
| Figure 1H | opto. silencing | *b1-GAL4 > GtACR1* | *w ; UAS-GtACR1 (attP40)/+ ; R71D08-p65ADZp (attP2) , R33E02-ZpGAL4DBD (VK00027)/+* |
| Figure 2A–C | opto. activation (control) | *empty* | *20XUAS-CsChrimson-mVenus (attP18)/+ ; R24A03-p65ADZp (attP40)/+ ; R74C01-ZpGdbd (attP2)/+* |
| Figure 2A–C | opto. activation | *b1* | *20XUAS-CsChrimson-mVenus (attP18)/+ ; +/+ ; R71D08-p65ADZp (attP2) , R33E02-ZpGAL4DBD (VK00027)/+* |
| Figure 2A–C | opto. activation | *b2* | *20XUAS-CsChrimson-mVenus (attP18)/+ ; VT40232-p65ADZp (attP40)/+ ; VT23830-ZpGAL4DBD (attP2)/+* |
| Figure 2D–F | opto. silencing (control) | *empty* | *w ; UAS-GtACR1 (attP40)/R24A03-p65ADZp (attP40) ; R74C01-ZpGdbd (attP2)/+* |
| Figure 2D–F | opto. silencing | *b1* | *w ; UAS-GtACR1 (attP40)/+ ; R71D08-p65ADZp (attP2) , R33E02-ZpGAL4DBD (VK00027)/+* |
| Figure 2D–F | opto. silencing | *b2* | *w ; UAS-GtACR1 (attP40)/VT40232-p65ADZp (attP40) ; VT23830-ZpGAL4DBD (attP2)/+* |
| Figure 3C,D | chronic silencing and pitch pert. | *b1-silenced or b1/Kir* | *w ; pJFRC79-8XLexAop2-FlpL (attP40)/+ ; pJFRC56-10XUAS-FRT > STOP > FRT-kir2.1-gfp (attP2)/R71D08-p65ADZp (attP2) , R33E02-ZpGAL4DBD (VK00027)* |
| Figure 3C,D | chronic silencing and pitch pert. (control) | *genetic control or b1/+* | *w ; +/+ ; R71D08-p65ADZp (attP2) , R33E02-ZpGAL4DBD (VK00027)/+* |
| Figure 3D | chronic silencing and pitch pert. (control) | *+/Kir* | *w ; pJFRC79-8XLexAop2-FlpL (attP40)/+ ; pJFRC56-10XUAS-FRT > STOP > FRT-kir2.1-gfp (attP2)/+* |
| Figure 3E,F | chronic silencing and pitch pert. | *b2-silenced or b2/Kir* | *w ; VT40232-p65ADZp (attP40)/+ ; pJFRC49-5XUAS-IVS-eGFPKir2.1 (attP2)/VT23830-ZpGAL4DBD (attP2)* |
| Figure 3E | chronic silencing and pitch pert. (control) | *b2/+* | *w ; VT40232-p65ADZp (attP40)/+ ; VT23830-ZpGAL4DBD (attP2)/+* |
| Figure 3E,F | chronic silencing and pitch pert. (control) | *genetic control or +/Kir* | *w ; +/+ ; pJFRC49-5XUAS-IVS-eGFPKir2.1 (attP2)/+* |
| Figure 3H,I | opto. silencing and pitch pert. (control) | *genetic control or empty* | *w ; UAS-GtACR1 (attP40)/R24A03-p65ADZp (attP40) ; R74C01-ZpGdbd (attP2)/+* |
| Figure 3H,I | opto. silencing and pitch pert. | *b1-silenced or b1* | *w ; UAS-GtACR1 (attP40)/+ ; R71D08-p65ADZp (attP2) , R33E02-ZpGAL4DBD (VK00027)/+* |
| Figure 3H,I | opto. silencing and pitch pert. | *b2-silenced or b2* | *w ; UAS-GtACR1 (attP40)/VT40232-p65ADZp (attP40) ; VT23830-ZpGAL4DBD (attP2)/+* |

**Table S3.** Full genotype of flies used in experiments (SI figures).

| Data shown in | Experiment type | Abbreviated genotype | Full genotype |
| --- | --- | --- | --- |
| Figure S1A,B | CNS anatomy | *b1-GAL4* | *20XUAS-CsChrimson-mVenus (attP18)/+ ; +/+ ; R71D08-p65ADZp (attP2) , R33E02-ZpGAL4DBD (VK00027)/+* |
| Figure S1C,D | CNS anatomy | *b2-GAL4* | *20XUAS-CsChrimson-mVenus (attP18)/+ ; VT40232-p65ADZp (attP40)/+ ; VT23830-ZpGAL4DBD (attP2)/+* |
| Figure S1E,F | phalloidin anatomy | *b1-GAL4* | *w ; 10XUAS-IVS-mCD8::GFP (attP40)/+ ; R71D08-p65ADZp (attP2) , R33E02-ZpGAL4DBD (VK00027)/+* |
| Figure S2A,C | opto. activation (control) | *empty* | *20XUAS-CsChrimson-mVenus (attP18)/+ ; R24A03-p65ADZp (attP40)/+ ; R74C01-ZpGdbd (attP2)/+* |
| Figure S2A,C | opto. activation | *b1* | *20XUAS-CsChrimson-mVenus (attP18)/+ ; +/+ ; R71D08-p65ADZp (attP2) , R33E02-ZpGAL4DBD (VK00027)/+* |
| Figure S2A,C | opto. activation | *b2* | *20XUAS-CsChrimson-mVenus (attP18)/+ ; VT40232-p65ADZp (attP40)/+ ; VT23830-ZpGAL4DBD (attP2)/+* |
| Figure S2A,C | opto. activation | *SS04528* | *20XUAS-CsChrimson-mVenus (attP18)/+ ; R19G08-p65ADZp (attP40)/+ ; R47F01-ZpGDBD (attP2)/+* |
| Figure S2B,D | opto. silencing (control) | *empty* | *w ; UAS-GtACR1 (attP40)/R24A03-p65ADZp (attP40) ; R74C01-ZpGdbd (attP2)/+* |
| Figure S2B,D | opto. silencing | *b1* | *w ; UAS-GtACR1 (attP40)/+ ; R71D08-p65ADZp (attP2) , R33E02-ZpGAL4DBD (VK00027)/+* |
| Figure S2B,D | opto. silencing | *b2* | *w ; UAS-GtACR1 (attP40)/VT40232-p65ADZp (attP40) ; VT23830-ZpGAL4DBD (attP2)/+* |
| Figure S2B,D | opto. silencing | *SS04528* | *w ; UAS-GtACR1 (attP40)/R19G08-p65ADZp (attP40) ; R47F01-ZpGDBD (attP2)/+* |
| Figure S3C,D | chronic silencing and roll pert. | *b1-silenced* or *b1/Kir* | *w ; pJFRC79-8XLexAop2-FlpL (attP40)/+ ; pJFRC56-10XUAS-FRT > STOP > FRT-kir2.1-gfp (attP2)/R71D08-p65ADZp (attP2) , R33E02-ZpGAL4DBD (VK00027)* |
| Figure S3C,D | chronic silencing and roll pert. (control) | *genetic control* or *b1/+* | *w ; +/+ ; R71D08-p65ADZp (attP2) , R33E02-ZpGAL4DBD (VK00027)/+* |
| Figure S3D | chronic silencing and roll pert. (control) | *+/Kir* | *w ; pJFRC79-8XLexAop2-FlpL (attP40)/+ ; pJFRC56-10XUAS-FRT > STOP > FRT-kir2.1-gfp (attP2)/+* |
| Figure S3E,F | chronic silencing and roll pert. | *b2-silenced* or *b2/Kir* | *w ; VT40232-p65ADZp (attP40)/+ ; pJFRC49-5XUAS-IVS-eGFPKir2.1 (attP2)/VT23830-ZpGAL4DBD (attP2)* |
| Figure S3E,F | chronic silencing and roll pert. (control) | *genetic control* or *b2/+* | *w ; VT40232-p65ADZp (attP40)/+ ; VT23830-ZpGAL4DBD (attP2)/+* |
| Figure S3F | chronic silencing and roll pert. (control) | *+/Kir* | *w ; +/+ ; pJFRC49-5XUAS-IVS-eGFPKir2.1 (attP2)/+* |
| Figure S4 | muscle imaging | *b1-silenced* or *b1/Kir* | *w ; 13XLexAop2-IVS-GCaMP6f-p10 (su(Hw)attP5)/R39E01-LexA (attP40) ; pJFRC49-10XUAS-IVS-eGFPKir2.1 (attP2)/R71D08-p65ADZp (attP2) , R33E02-ZpGAL4DBD (VK00027)* |
| Figure S4 | muscle imaging | *genetic control* or *b1/+* | *w ; 13XLexAop2-IVS-GCaMP6f-p10 (su(Hw)attP5)/R39E01-LexA (attP40) ; R71D08-p65ADZp (attP2) , R33E02-ZpGAL4DBD (VK00027)/+* |
| Figure S4 | muscle imaging | *genetic control* or *+/Kir* | *w ; 13XLexAop2-IVS-GCaMP6f-p10 (su(Hw)attP5)/R39E01-LexA (attP40) ; pJFRC49-10XUAS-IVS-eGFPKir2.1 (attP2)/+* |
| Figure S5A,B | anatomy | *SS04528* | *20XUAS-CsChrimson-mVenus (attP18)/+ ; R19G08-p65ADZp (attP40)/+ ; R47F01-ZpGDBD (attP2)/+* |
| Figure S5C–E | opto. activation (control) | *empty* | *20XUAS-CsChrimson-mVenus (attP18)/+ ; R24A03-p65ADZp (attP40)/+ ; R74C01-ZpGdbd (attP2)/+* |
| Figure S5C–E | opto. activation | *b1* | *20XUAS-CsChrimson-mVenus (attP18)/+ ; +/+ ; R71D08-p65ADZp (attP2) , R33E02-ZpGAL4DBD (VK00027)/+* |
| Figure S5C–E | opto. activation | *SS04528* | *20XUAS-CsChrimson-mVenus (attP18)/+ ; R19G08-p65ADZp (attP40)/+ ; R47F01-ZpGDBD (attP2)/+* |
| Figure S5F–H | opto. silencing (control) | *empty* | *w ; UAS-GtACR1 (attP40)/R24A03-p65ADZp (attP40) ; R74C01-ZpGdbd (attP2)/+* |
| Figure S5F–H | opto. silencing | *b1* | *w ; UAS-GtACR1 (attP40)/+ ; R71D08-p65ADZp (attP2) , R33E02-ZpGAL4DBD (VK00027)/+* |
| Figure S5F–H | opto. silencing | *SS04528* | *w ; UAS-GtACR1 (attP40)/R19G08-p65ADZp (attP40) ; R47F01-ZpGDBD (attP2)/+* |
| Figure S6A | chronic silencing and pitch pert. | *b1/Kir* | *w ; +/+ ; pJFRC49-5XUAS-IVS-eGFPKir2.1 (attP2)/R71D08-p65ADZp (attP2) , R33E02-ZpGAL4DBD (VK00027)* |
| Figure S6A | chronic silencing and pitch pert. (control) | *b1/+* | *w ; +/+ ; R71D08-p65ADZp (attP2) , R33E02-ZpGAL4DBD (VK00027)/+* |
| Figure S6A | chronic silencing and pitch pert. | *b2/Kir* | *w ; VT40232-p65ADZp (attP40)/+ ; pJFRC49-5XUAS-IVS-eGFPKir2.1 (attP2)/VT23830-ZpGAL4DBD (attP2)* |
| Figure S6A | chronic silencing and pitch pert. (control) | *b2/+* | *w ; VT40232-p65ADZp (attP40)/+ ; VT23830-ZpGAL4DBD (attP2)/+* |
| Figure S6A | chronic silencing and pitch pert. (control) | *+/Kir* | *w ; +/+ ; pJFRC49-5XUAS-IVS-eGFPKir2.1 (attP2)/+* |
| Figure S6B | chronic silencing and roll pert. | *b1/Kir* | *w ; +/+ ; pJFRC49-5XUAS-IVS-eGFPKir2.1 (attP2)/R71D08-p65ADZp (attP2) , R33E02-ZpGAL4DBD (VK00027)* |
| Figure S6B | chronic silencing and roll pert. (control) | *b1/+* | *w ; +/+ ; R71D08-p65ADZp (attP2) , R33E02-ZpGAL4DBD (VK00027)/+* |
| Figure S6B | chronic silencing and roll pert. | *b2/Kir* | *w ; VT40232-p65ADZp (attP40)/+ ; pJFRC49-5XUAS-IVS-eGFPKir2.1 (attP2)/VT23830-ZpGAL4DBD (attP2)* |
| Figure S6B | chronic silencing and roll pert. (control) | *b2/+* | *w ; VT40232-p65ADZp (attP40)/+ ; VT23830-ZpGAL4DBD (attP2)/+* |
| Figure S6B | chronic silencing and roll pert. (control) | *+/Kir* | *w ; +/+ ; pJFRC49-5XUAS-IVS-eGFPKir2.1 (attP2)/+* |
| Figure S7A | opto. silencing and pitch pert. | *b1* | *w ; UAS-GtACR1 (attP40)/+ ; R71D08-p65ADZp (attP2) , R33E02-ZpGAL4DBD (VK00027)/+* |
| Figure S7A | opto. silencing and pitch pert. | *SS04528* | *w ; UAS-GtACR1 (attP40)/R19G08-p65ADZp (attP40) ; R47F01-ZpGDBD (attP2)/+* |
| Figure S7A | opto. silencing and pitch pert. | *b2* | *w ; UAS-GtACR1 (attP40)/VT40232-p65ADZp (attP40) ; VT23830-ZpGAL4DBD (attP2)/+* |
| Figure S7A | opto. silencing and pitch pert. (control) | *empty* | *w ; UAS-GtACR1 (attP40)/R24A03-p65ADZp (attP40) ; R74C01-ZpGdbd (attP2)/+* |
| Figure S7B | opto. silencing and roll pert. | *b1* | *w ; UAS-GtACR1 (attP40)/+ ; R71D08-p65ADZp (attP2) , R33E02-ZpGAL4DBD (VK00027)/+* |
| Figure S7B | opto. silencing and roll pert. | *SS04528* | *w ; UAS-GtACR1 (attP40)/R19G08-p65ADZp (attP40) ; R47F01-ZpGDBD (attP2)/+* |
| Figure S7B | opto. silencing and roll pert. | *b2* | *w ; UAS-GtACR1 (attP40)/VT40232-p65ADZp (attP40) ; VT23830-ZpGAL4DBD (attP2)/+* |
| Figure S7B | opto. silencing and roll pert. (control) | *empty* | *w ; UAS-GtACR1 (attP40)/R24A03-p65ADZp (attP40) ; R74C01-ZpGdbd (attP2)/+* |
| Figure S8A | chronic silencing and pitch pert. (control) | *genetic control* | *w ; +/+ ; R71D08-p65ADZp (attP2) , R33E02-ZpGAL4DBD (VK00027)/+* |
| Figure S8B | chronic silencing and pitch pert. | *b1-silenced* | *w ; pJFRC79-8XLexAop2-FlpL (attP40)/+ ; pJFRC56-10XUAS-FRT > STOP > FRT-kir2.1-gfp (attP2)/R71D08-p65ADZp (attP2) , R33E02-ZpGAL4DBD (VK00027)* |
| Figure S8C | chronic silencing and pitch pert. (control) | *genetic control* | *w ; +/+ ; pJFRC49-5XUAS-IVS-eGFPKir2.1 (attP2)/+* |
| Figure S8D | chronic silencing and pitch pert. | *b2-silenced* | *w ; VT40232-p65ADZp (attP40)/+ ; pJFRC49-5XUAS-IVS-eGFPKir2.1 (attP2)/VT23830-ZpGAL4DBD (attP2)* |
| Figure S9A,B | opto. silencing and pitch pert. (control) | *empty* | *w ; UAS-GtACR1 (attP40)/R24A03-p65ADZp (attP40) ; R74C01-ZpGdbd (attP2)/+* |
| Figure S9A,B | opto. silencing and pitch pert. | *b1* | *w ; UAS-GtACR1 (attP40)/+ ; R71D08-p65ADZp (attP2) , R33E02-ZpGAL4DBD (VK00027)/+* |
| Figure S9A,B | opto. silencing and pitch pert. | *b2* | *w ; UAS-GtACR1 (attP40)/VT40232-p65ADZp (attP40) ; VT23830-ZpGAL4DBD (attP2)/+* |

**Table S4.** Model parameters

| Symbol | Definition | Value |
| --- | --- | --- |
| $m$ | body mass | 1.34 mg<sup>37</sup> |
| $I_{\text{pitch}}$ | pitch moment of inertia | 0.506 mg mm<sup>2</sup><sup>25</sup> |
| $R$ | wing span length | 2.13 mm |
| $\bar{c}$ | mean wing chord length | 0.7 mm |
| $S$ | wing area | 5.74 mm<sup>2</sup><sup>37</sup> |
| $\hat{r}_2^2(S)$ | non-dimensionalized second moment of wing area | 0.313<sup>25</sup> |
| $f$ | wingbeat frequency | 225 Hz |
| $\phi_0$ | stroke angle offset | 97.3° |
| $\phi_m$ | stroke angle amplitude | 67.6° |
| $K_\phi$ | stroke angle waveform parameter | 0.7<sup>9</sup> |
| $\theta_0$ | deviation angle offset | 0° |
| $\theta_m$ | deviation angle amplitude | 0° |
| $\delta_\theta$ | deviation angle phase shift | 0° |
| $\eta_0$ | rotation angle offset | 90.0° |
| $\eta_m$ | rotation angle amplitude | 67.3° |
| $\delta_\eta$ | rotation angle phase shift | -85.0° |
| $C_\eta$ | rotation angle waveform parameter | 2.4<sup>9</sup> |
| $\rho$ | air density | 1.2 kg m<sup>-3</sup> |
| $g$ | gravitational acceleration | 9.8 m s<sup>-2</sup> |
| $C_{\text{friction}}$ | body pitch rotational drag | 0.52 mg mm<sup>2</sup><sup>25</sup> |
| $C_{\text{rot}}$ | rotational force coefficient | 1.57<sup>11</sup> |
| $C_{L_{\text{max}}}$ | maximum lift coefficient | 1.8<sup>15,18</sup> |
| $C_{D_{\text{max}}}$ | maximum drag coefficient | 3.4<sup>15,18</sup> |
| $C_{D_0}$ | minimum drag coefficient | 0.4<sup>15,18</sup> |
| $T_{\text{pulse}}$ | magnetic perturbation duration | 7 ms or 15 ms |
| $\theta_{\text{pin}}$ | angle between body and pin | 45° |
| $r_{\text{hinge}}$ | distance from body cm to wing hinge | 0.22 mm |

## Supplementary Movies

**Movie S1. Optogenetic excitation of a b1-GAL4 > CsChrimson fly**. High speed video footage of a *b1-GAL4 > CsChrimson* fly undergoing a 50 ms bout of red light stimulus (indicated by red square in corner of screen). The three panels show two side views (left and right) and one top view (middle). Panel in bottom left corner indicates time in the recording relative to the onset of light stimulus (*t*=0 ms), measured in milliseconds. This video corresponds to the photomontage labeled “1” in Figure 1G. If not attached, the video can be found at https://youtu.be/6auUKQ_MxpM.

**Movie S2. Optogenetic excitation of a b1-GAL4 > CsChrimson fly (second example)**. A second example of a *b1-GAL4 > CsChrimson* fly undergoing a optogenetic stimulation, corresponding to the photomontage labeled “2” in Figure 1G. If not attached, the video can be found at https://youtu.be/fdlSEWqpatk.

**Movie S3. Optogenetic silencing of a b1-GAL4 > GtACR1 fly**. High speed video footage of a *b1-GAL4 > GtACR1* fly undergoing a 50 ms bout of green light stimulus (indicated by green square in corner of screen). This video corresponds to the photomontage labeled “1” in Figure 1H. If not attached, the video can be found at https://youtu.be/nQRCrrqovxw.

**Movie S4. Optogenetic silencing of a b1-GAL4 > GtACR1 fly (second example)**.A second example of a *b1-GAL4 > GtACR1* fly undergoing undergoing optogenetic inhibition, correspondng to the photomontage labeled “2” in Figure 1H. If not attached, the video can be found at https://youtu.be/IfPbtp84R5k.

**Movie S5. Optogenetic excitation of 24 b1-GAL4 > CsChrimson flies**. Side views from 24 movies showing *b1-GAL4 > CsChrimson* flies undergoing a 50 ms bout of red light stimulus (indicated by red square in corner of screen). If not attached, the video can be found at https://youtu.be/Ea8QCYYBC6U.

**Movie S6. Optogenetic excitation of 24 b2-GAL4 > CsChrimson flies**. Side views from 24 movies showing *b2-GAL4 > CsChrimson* flies undergoing a 50 ms bout of red light stimulus (indicated by red square in corner of screen). If not attached, the video can be found at https://youtu.be/wDalFmkic7g.

**Movie S7. Optogenetic excitation of 24 SS01062-GAL4 > CsChrimson flies**. Side views from 24 movies showing *SS01062-GAL4 > CsChrimson* flies (genetic control) undergoing a 50 ms bout of red light stimulus (indicated by red square in corner of screen). If not attached, the video can be found at https://youtu.be/1da_1aPYARs.

**Movie S8. Optogenetic inhibition of 24 b1-GAL4 > GtACR1 > flies**. Side views from 24 movies showing *b1-GAL4 > GtACR1* flies undergoing a 50 ms bout of green light stimulus (indicated by green square in corner of screen). If not attached, the video can be found at https://youtu.be/LpOOnIQ9qpk.

**Movie S9. Optogenetic inhibition of 24 b2-GAL4 > GtACR1 flies**. Side views from 24 movies showing *b2-GAL4 > GtACR1* undergoing a 50 ms bout of green light stimulus (indicated by green square in corner of screen). If not attached, the video can be found at https://youtu.be/f1xQ0I48S0E.

**Movie S10. Optogenetic inhibition of 24 SS01062-GAL4 > GtACR1 flies**. Side views from 24 movies showing *SS01062-GAL4 > GtACR1* flies (genetic control) undergoing a 50 ms bout of green light stimulus (indicated by green square in corner of screen). If not attached, the video can be found at https://youtu.be/49Y7GfolqHE.

**Movie S11. Video data for pitch perturbation of genetic control group fly in Figure 3C**. Video footage showing two side views (left and right) and one overhead view (middle) of a genetic control fly undergoing a 7 ms magnetic pitch perturbation (indicated by yellow square the corner of each view). Bottom left corner shows time, measured in milliseconds, relative to the onset of the magnetic pulse at *t*=0 ms. If not attached, the video can be found at https://youtu.be/Mt3IQCYg644.

**Movie S12. Video data for pitch perturbation of b1 motoneuron silenced fly in Figure 3C**. Video footage showing two side views (left and right) and one overhead view (middle) of a b1-silenced fly undergoing a 7 ms magnetic pitch perturbation, as in Movie S11. If not attached, the video can be found at https://youtu.be/KW12ydQGpp4.

**Movie S13. Video data for pitch perturbation of genetic control group fly in Figure 3E**. Video footage showing two side views (left and right) and one overhead view (middle) of a genetic control fly undergoing a 7 ms magnetic pitch perturbation, as in Movies S11 and S12. If not attached, the video can be found at https://youtu.be/acw9N-NOFfI.

**Movie S14. Video data for pitch perturbation of b2 motoneuron silenced fly in Figure 3C**. Video footage showing two side views (left and right) and one overhead view (middle) of a b2-silenced fly undergoing a 7 ms magnetic pitch perturbation, as in Movies S11 and S12. If not attached, the video can be found at https://youtu.be/7B6LzRhje_0.

## Footnotes

1 Note that the author who tuned *B* was blind to the range of values fitted in the nonlinear simulations.

## Notes

### Competing Interest Statement

The authors have declared no competing interest.

### Summary of Updates

Adding middle initial to one author name

